# SCARF1-induced efferocytosis plays an immunomodulatory role in humans, and autoantibodies targeting SCARF1 are produced in patients with systemic lupus erythematosus

**DOI:** 10.1101/2021.05.24.445434

**Authors:** April M. Jorge, Taotao Lao, Rachel Kim, Samantha Licciardi, Joseph ElKhoury, Andrew Luster, Terry K. Means, Zaida G. Ramirez-Ortiz

**Affiliations:** Clinical Epidemiology Program of the Division of Rheumatology, Allergy, and Immunology and Mongan Institute, Department of Medicine, Massachusetts General Hospital, 100 Cambridge Street, Suite 1600, Boston, MA 02114; Rheumatology Unit, Division of Rheumatology, Allergy, and Immunology, Massachusetts General Hospital, 55 Fruit Street, Yawkey 2C, Boston, MA 02114; Center for Immunology and Inflammatory Diseases. Department of Rheumatology, Allergy and Immunology. Massachusetts General Hospital. 13st CNY 149, Charlestown MA 02129; Sanofi, Autoimmunity Cluster, Immunology & Inflammation Research Therapeutic Area. 270 Albany Street Room 2427-A, Cambridge MA 02139; Department of Medicine, Division of Infectious Diseases and Immunology. University of Massachusetts Medical School. 364 Plantation St. LRB319, Worcester MA 01605

**Keywords:** Scavenger Receptors, SCARF1, Lupus, efferocytosis, IL-10

## Abstract

Deficiency in the clearance of cellular debris is a major pathogenic factor in the emergence of autoimmune diseases. We previously demonstrated that mice deficient for scavenger receptor class F member 1 (SCARF1) develop a lupus-like autoimmune disease with symptoms similar to human systemic lupus erythematosus (SLE), including a pronounced accumulation of apoptotic cells (ACs). Therefore, we hypothesized that SCARF1 will be important for clearance of ACs and maintenance of self-tolerance *in humans*, and that dysregulation of this process *could* contribute to SLE. Here, we show that SCARF1 is highly expressed on phagocytic cells, where it functions as an efferocytosis receptor. In healthy individuals, we discovered that engagement of SCARF1 by ACs on BDCA1^+^ dendritic cells (DCs) initiates an interleukin-10 (IL-10) anti-inflammatory response mediated by the phosphorylation of signal transducer and activator of transcription 1 (STAT1). Unexpectedly, there was no significant difference in SCARF1 expression in SLE patient samples compared to healthy donor samples. However, we detected anti-SCARF1 autoantibodies in 26% of SLE patients, which was associated with dsDNA antibody positivity. Furthermore, our data shows a direct correlation of the levels of anti-SCARF1 in the serum and defects in the removal of ACs. Depletion of immunoglobulin restores efferocytosis in SLE serum, suggesting that defects in the removal of ACs is partially mediated by SCARF1 pathogenic autoantibodies. Our data demonstrate that human SCARF1 is an AC receptor in DCs and plays a role in maintaining tolerance and homeostasis.

## Introduction

The human body generates millions of cells daily, and the same number of cells must die to maintain homeostasis in the body (1). In a healthy individual, apoptotic cells (ACs) are efficiently removed before debris accumulates, avoiding an inflammatory response (2). However, inefficient clearance of ACs can result in the accumulation of apoptotic debris, leading to a break in tolerance and development of autoimmunity (3). Indeed, patients with systemic lupus erythematosus (SLE) have increased levels of circulating ACs, indicating a failure in the clearance of dying cells (4, 5). Uncleared ACs can undergo secondary necrosis and can accumulate in germinal centers, where they can activate complement and autoreactive B cells. Furthermore, noxious intracellular molecules are released from secondary necrotic cells resulting in the production of autoantibodies, a hallmark feature of lupus (6).

The controlled elimination of dying cells is initiated when so-called death receptors interact with their cognate ligands (2). During apoptosis, phosphatidylserine is externalized from the inner leaflet of the cell membrane, where it serves as the primary “eat me” signal for phagocytes (7). Several receptors (TIM-3, TAM) or soluble bridging proteins (C1q, calreticulin and milk-fat globule epidermal growth factor 8 [MFG-E8]) specifically bind to phosphatidylserine exposed on the surface of ACs to enhance the uptake and rapid removal by phagocytes (8–11). Studies have demonstrated an essential role for C1q in AC clearance and the development of autoimmunity, but these mechanisms require further characterization (8, 12). Therefore, additional studies are necessary to understand how phagocytes capture and engulf ACs, as well as the signaling pathways initiated by cellular debris to prevent the loss of tolerance. This lack of knowledge remains a critical barrier to understanding autoimmune disease pathogenesis, especially in SLE.

We previously demonstrated that the scavenger receptor SCARF1 (scavenger receptor class F member 1, also known as SR-F1 or SREC1) is a non-redundant AC receptor in mice (13). SCARF1 belongs to the scavenger receptor (SR) superfamily of proteins that is defined by their ability to bind and endocytose a wide range of ligands (14). The SR family was originally identified as modified low-density lipoprotein receptors, but, over the last two decades, new SR members have been identified (15). SRs are divided into different classes (A-J), sharing little or no structural homology (14, 16). SCARF1 is an 86 KDa type-I transmembrane protein composed of an extracellular region with several epidermal growth factor (EGF)-like domains, a short cytoplasmic region, and a long cytoplasmic tail that is serine and proline-rich (17). In our previous study, we demonstrated that mice with global *Scarf1* deficiency spontaneously develop autoimmune disease with clinical manifestations that are strikingly similar to human SLE and have pronounced accumulation of AC (13). However, the role of human SCARF1 in the removal of ACs is unknown.

Based on our data from *Scarf1* deficient mice, we hypothesized that SCARF1 mediates AC clearance in humans and dysregulation of SCARF1 in SLE patients results in the accumulation of ACs and contributes to SLE disease. To date, only one study has characterized the cellular distribution of SCARF1 in humans during disease. Patten *et al*. showed that SCARF1 contributes to lymphocyte adhesion to hepatic sinusoidal endothelial cells, where they could play a role in the inflammatory response (18). Thus, studies are needed to understand human SCARF1 biology and how this SR contributes to disease.

In this study, we investigated the role of human SCARF1 as an efferocytosis receptor. We demonstrate the presence of SCARF1 in immune phagocytes. In BDCA1^+^ myeloid DCs (CD1c^+^ CD11c^+^ DCs), SCARF1 is responsible for the binding and phagocytosis of ACs. Furthermore, we observed that exposing SCARF1^+^BDCA1^+^ DCs to ACs results in the upregulation of SCARF1, IL-10 and the phosphorylation of STAT1. Finally, we did not observe a significant difference in membrane-bound SCARF1 between lupus patients and healthy controls. However, we did observe an increase in autoantibodies to SCARF1 in the serum of SLE patients, indicating that antibodies to SCARF1 might contribute to the breakdown of self-tolerance by impairing the removal of ACs in some patients. Together, these data suggest a role for SCARF1 in the pathogenesis of autoimmune diseases, particularly SLE.

## Results

### SCARF1 is expressed on phagocytic cells and recognizes apoptotic cells

We previously showed that SCARF1 is an AC receptor expressed on CD8a^+^ DCs in a mouse model; however, in humans, the expression and function of SCARF1 remains unknown. We wanted to test the capacity of SCARF1 in the removal of ACs in a human cell system. Endothelial cells express high levels of SCARF1 (17, 19); therefore, we decided to use a human endothelial cell line to answer these questions. We successfully eliminated SCARF1 from an endothelial cell line (a telomerase-immortalized human microvascular endothelium cell line, TIME) using CRISPR-Cas9 (Fig. 1A). In the absence of SCARF1, we observed a significant reduction in the uptake of ACs (Fig. 1B). Similar to our previous findings using the *Scarf1*^-/-^ mouse model, we observed a 20–40% reduction in the removal of ACs in SCARF1-deficient cells. These findings suggest that SCARF1 expressed on human cells mediates AC binding and uptake.

**Figure 1.**
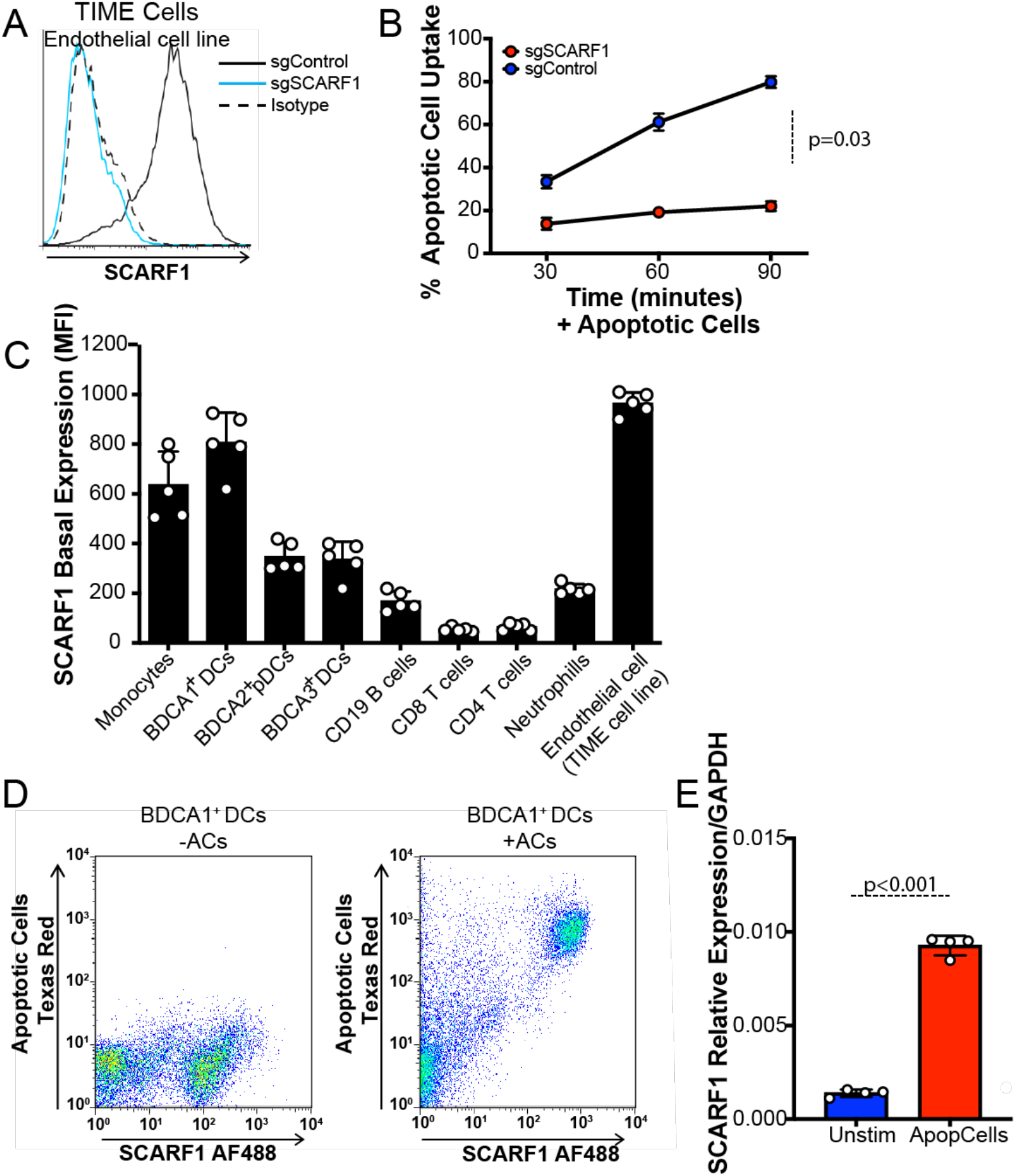
SCARF1 profiling in immune cells. (A) SCARF1 is highly expressed on endothelial cells. CRISPR-*Cas9* deletion of SCARF1 was performed on the human endothelial cell line, TIME. The histogram is representative of three independent experiments. (B) Uptake of apoptotic cells in SCARF1^+^ (sgControl) and SCARF1^-^ (sgSCARF1) endothelial cells. Cells were stimulated with apoptotic cells for 30, 60, or 90 minutes. Apoptotic cell uptake was measured by flow cytometry. Results shown are the average percent uptake ±SD of two independent experiments performed in triplicates. *P*=0.03 by Student’s *t*-test analysis. (C) SCARF1 is expressed on phagocytes. PBMCs were isolated from the blood of healthy donors (n=5) and stained for flow cytometry against SCARF1 and immune cell markers. TIME cells were used as a positive control. MFI, mean fluorescence intensity. Data represent the mean (±SEM) of two independent experiments with 2–3 replicates. (D) Representative flow cytometry plot of total SCARF1 expression in the presence or absence of apoptotic cells. Data shown are representative of three independent experiments with similar results. (E) SCARF1 is upregulated after stimulation with apoptotic cells. BDCA1^+^ DCs were stimulated with apoptotic cells for 4 hours. Cells were lysed, and mRNA was measured using qPCR. Data represent the mean (±SEM) of two independent experiments in duplicate. *P*<0.001

To characterize relevant SCARF1-expressing cells in the human immune system, we used flow cytometry to measure the basal levels of SCARF1 in peripheral blood mononuclear cells (PBMCs) from healthy donors (n=5, Fig. 1C). We detected SCARF1 expression on phagocytic cells, like DCs and monocytes, but absent on T cells and lower on B cells, suggesting a role for SCARF1 in the recognition and/or removal of foreign particles. Our initial observations in the mouse system showed an important role for SCARF1 expressed on DCs for the removal of ACs. Previous work identified BDCA1^+^ DCs (CD1c^+^ DCs) and BDCA3^+^ (CD141^+^ DCs) in the removal and cross-presentation of necrotic cell antigens (20–22). However, little is known about the mechanism and function of BDCA1^+^ DCs in efferocytosis, the process by which phagocytes remove ACs. Therefore, we decided to investigate the role of SCARF1 in BDCA1^+^ DCs. Initially, we investigated the regulation of SCARF1 in the presence of ACs. We observed that SCARF1 interacts with apoptotic debris in BDCA1^+^ DCs by flow cytometry (Fig. 1D). In addition, we detected an upregulation of SCARF1 mRNA after stimulation with ACs (Fig. 1E). Our data indicate a novel role for SCARF1 expressed on BDCA1^+^ DCs in the recognition and uptake of ACs.

### SCARF1 is an efferocytosis receptor for apoptotic cells

We observed an interaction between SCARF1 and ACs; thus, we wanted to determine whether SCARF1 is a phagocytic receptor. First, we added ACs to PBMCs and then assayed for surface expression of SCARF1. The addition of ACs significantly reduced SCARF1 membrane expression in all immune phagocytes, with up to an 80% reduction in SCARF1 surface expression in monocytes. These data indicate that SCARF1 becomes internalized (Fig. 2A), supporting our hypothesis that SCARF1 mediates the binding and clearance of ACs.

**Figure 2.**
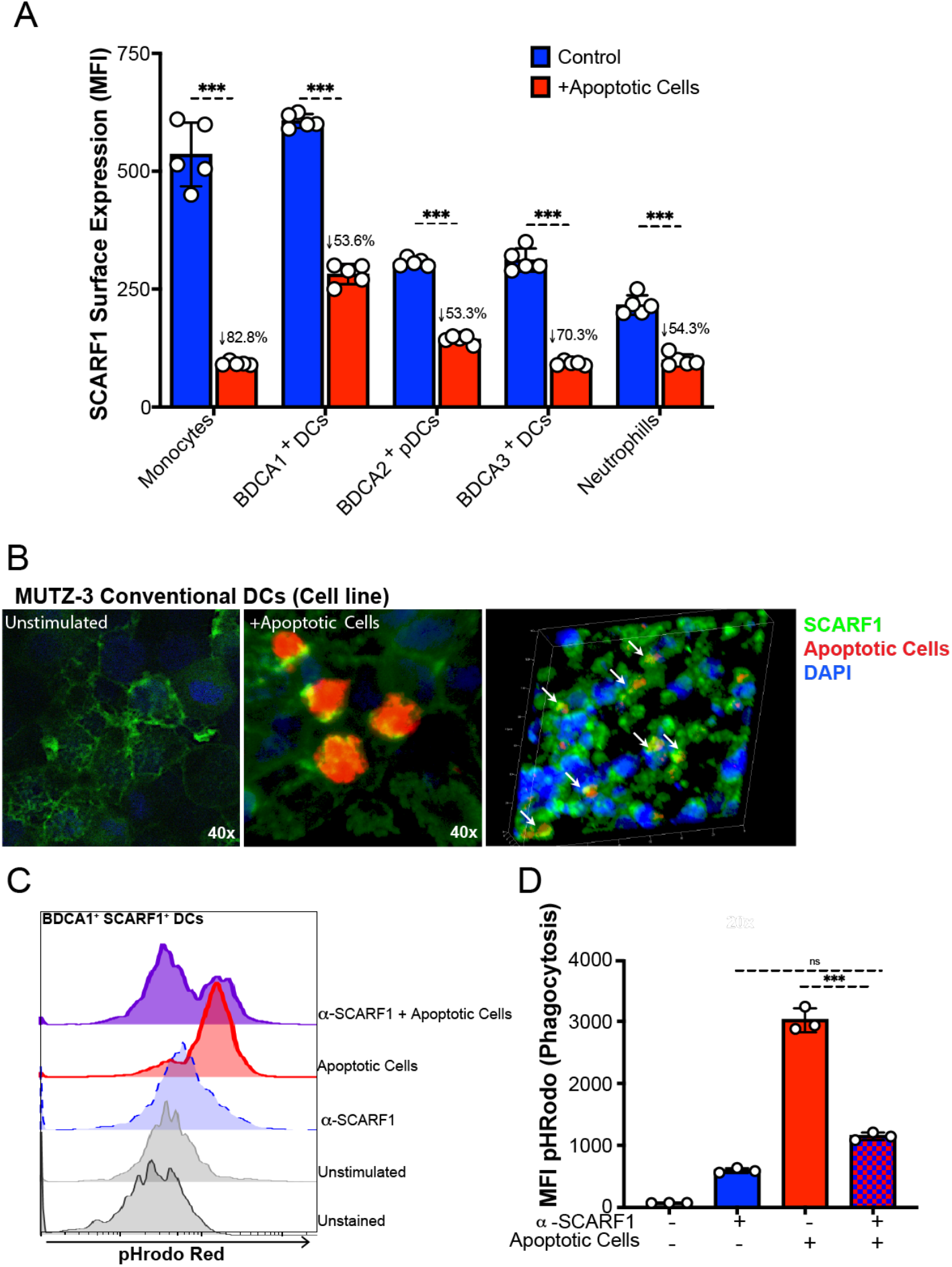
SCARF1 is an efferocytosis receptor on BDCA1^+^ DCs. (A) SCARF1 is internalized after interacting with apoptotic cells. PBMCs (1×10^6^ cells/mL) from healthy controls (n=5) were stimulated with apoptotic cells (2×10^6^ cells/mL) for 1 hour. After 1 hour, media was removed, and new media was added. Cells were incubated for 3 hours, and PBMCs were stained for flow cytometry. Data represent the mean (±SEM) of three independent experiments with 1–2 independent donors. Percent reduction is shown on the graph. \*\*\**P*<0.001. (B) SCARF1 is a phagocytic receptor for apoptotic cells in conventional DCs. MUTZ-3 DC line confocal images of SCARF1 (green), DNA (blue, DAPI), and apoptotic cells (red) 2 hours post-stimulation. Representative images from 2 independent experiments with 3 replicates. Scale=40x (left and middle), 20x (right). Arrows show the colocalization of apoptotic cells and SCARF1. (C-D) Blocking SCARF1 reduces apoptotic cell uptake. PBMCs (1×10^6^/mL) were incubated with anti-SCARF1 for 30 minutes, then stimulated with pHrodo red-labeled apoptotic cells (2×10^6^/mL). Cells were incubated for 3 hours and then stained and analyzed by flow cytometry. (C) Representative histogram of 3 independent experiments. (D) MFI quantification of internalization by pHrodo. Data represents the mean (±SEM) of 3 independent experiments, *P*< *(0.05) ***(0.0001) by two-way ANOVA.

To prove that SCARF1 gets internalized with the apoptotic debris, we performed confocal microscopy using the cell line MUTZ-3. MUTZ-3 is a human monocytic cell line derived from the peripheral blood of patients with acute myeloid leukemia (23). This cell line is capable of maturing into functional DCs, expresses co-stimulatory molecules, and is comparable to monocyte-derived DCs (24, 25). As previously shown, SCARF1 is expressed on the cell surface of naïve cells (26) (Fig. 2B, left panel). Upon stimulation with ACs, SCARF1 gets internalized (Fig. 2B, middle panel) and colocalizes with ACs (Fig. 2B, right panel, white arrows).

There is considerable functional redundancy in the receptors that mediate AC clearance (27). To demonstrate that SCARF1 is a non-redundant efferocytosis receptor, we blocked SCARF1 on BDCA1^+^ DCs by pretreating these cells with anti-human SCARF1 antibodies and investigated the phagocytosis of ACs. We labeled ACs with pHrodo red, which increases in fluorescence with low pH (i.e., after AC internalization), and assayed for phagocytosis by flow cytometry (Fig. 2C-D). Our results show a significant (*p*=0.0001) decrease in the removal of ACs in the SCARF1-blocked BDCA1^+^ DCs. We observed over 60% reduction in phagocytosis in SCARF1-blocked DCs, comparable to our prior observations in SCARF1-deficient mice (13). We previously reported that the complement protein C1q binds to ACs, leading to their uptake. However, using a human cell system, we did not see a significant difference in the presence or absence of C1q (data not shown). Together, these data suggest that SCARF1 is a non-redundant efferocytosis receptor on human BDCA1^+^ DCs.

### Efferocytosis by SCARF1+ BDCA1+ dendritic cells induce an immunomodulatory gene expression profile

We observed quick and efficient removal of ACs by SCARF1-expressing DCs. However, little is known about the SCARF1 pathway and what role it plays in the immune response. This made us wonder what other signatures were present during the efferocytosis program. To address these questions, we sorted SCARF1^Lo^ and SCARF1^Hi^ PBMCs (Supplemental Fig. 1A) to analyze the expression profile in defined subsets of BDCA1^+^ DCs that express or lack SCARF1. We are aware that the majority of the cells are intermediate for SCARF1 protein expression; however, we wanted to determine the contribution of SCARF1^Lo^ vs. SCARF1^Hi^ in the efferocytosis process. We used a Nanostring immunology panel to address these questions (Fig. 3A and Supplemental Fig. 1B).

**Figure 3.**
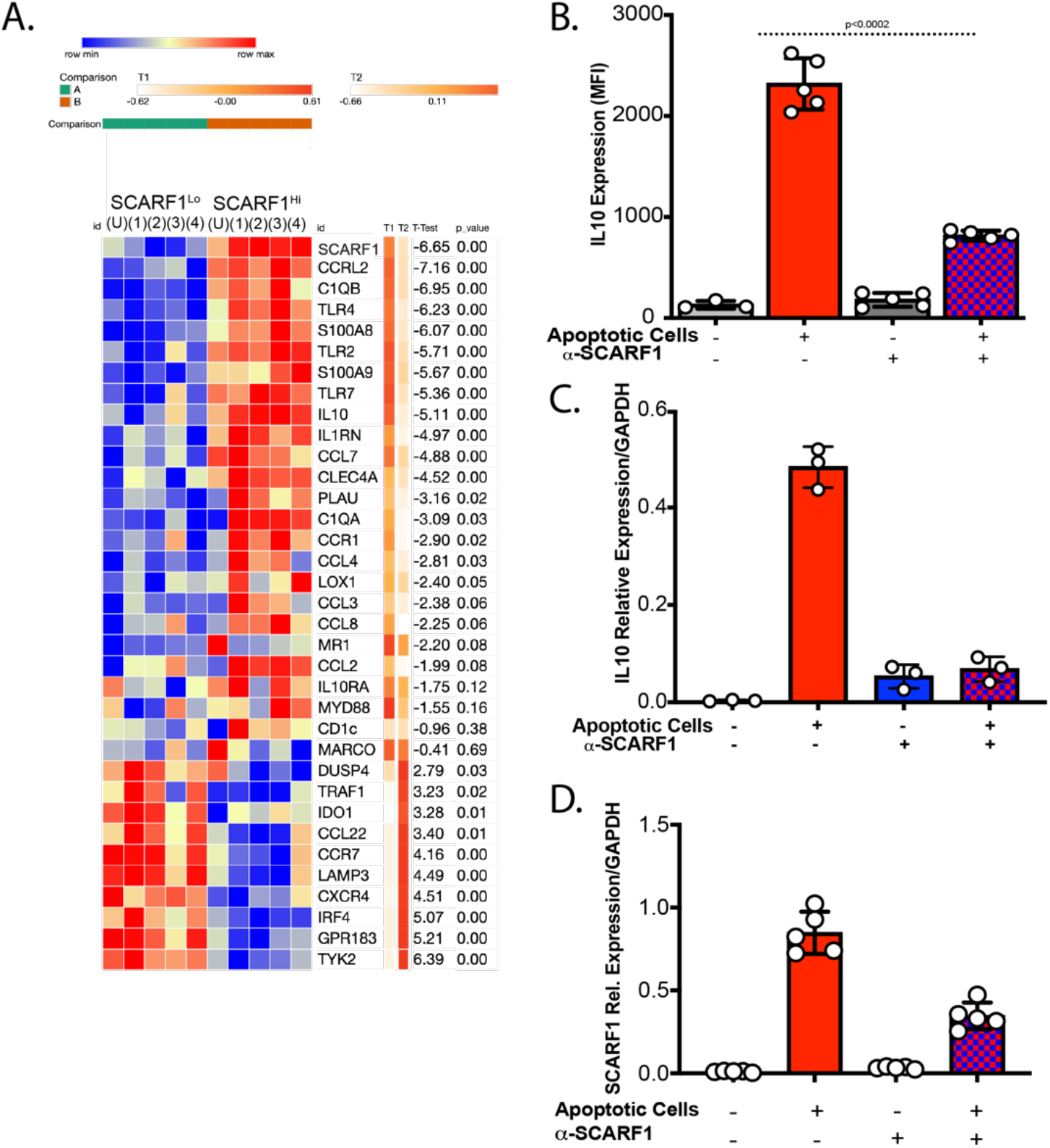
SCARF1 is involved in the inflammatory response. (A) Nanostring gene profiling of SCARF1^Lo^ and SCARF1^Hi^ BDCA1^+^ DCs. PBMCs were isolated from healthy controls and treated with apoptotic cells for 4 hours. Live BDCA1^+^ DCs (CD11c^+^BDCA1^+^Live/dead-UV) were sorted into SCARF1 negative (SCARF1^Lo^) or SCARF1 positive (SCARF1^Hi^). mRNA was extracted, and genetic profiling was performed using a Nanostring immunology panel. Heatmap showing the top 35 differentially expressed genes (log 2) after normalization using Morpheus software. Representative heatmap of single Nanostring assay (n=4). U, unstimulated. P values by Student’s *t*-test. (B-D) Expression of IL10 on BDCA1^+^ DCs. BDCA1^+^ sorted DCs (1×10^6^/mL) were block or not with anti-SCARF1 blocking antibody for 30 min, then they were stimulated with Live/Dead-labeled apoptotic cells (2×10^6^/mL) for 4 hours. (B) Cells were stained and analyzed for flow cytometry CD11c^+^CD11b^+^BDCA1^+^SCARF1^+^ and intracellular stain for IL10. MFI, mean fluorescent intensity. Data represent the mean (±SEM) of 3 independent experiments. *P*<0.002 by two-way ANOVA. (C) Cells were lysed, and mRNA was measured using qPCR. Data are the mean (±SEM) of 2 independent experiments with n=3. *P*=0.01 by Mann-Whitney test (D) Cells were lysed, and mRNA was measured using qPCR. Data are the mean (±SEM) of 2 independent experiments with n=5. *P*=0.003 by Mann-Whitney test.

We observed a clear differential expression of genes in SCARF1^Lo^ and SCARF1^Hi^ DCs (Fig. 3A, Supplemental Fig. 1B). SCARF1^Hi^ DCs express a variety of inflammatory genes, including chemokines, that are responsible for the proper recruitment of immune cells. We observed upregulation of chemokine (C-C motif) ligand (CCL)7, CCL8, CCL2, CCL4 and CCR1. These chemokines are immunomodulatory and are involved in the recruitment of monocytes and DCs (28) that are essential for the rapid removal of ACs. As previously described, we also observed the expression of Toll-like receptors (TLRs) in the presence of ACs and associated nucleic acids (29). Our data show the upregulation of TLR2, TLR4, and TLR7. Activation of these TLRs has been shown to modulate the presentation of cellular antigens (30). We observed a slight downregulation of scavenger receptors, macrophage receptor with collagenous structure (MARCO), and lectin-type oxidized LDL receptor 1 (LOX1), suggesting that receptors for danger signals are not involved in the recognition of ACs.

As expected, we did not observe the expression of type I interferon and the kinase tyrosine kinase 2 (Tyk2) is downregulated in our model, which is consistent with efferocytosis being an immunologically-silent process (31). Our data show the upregulation of the immunomodulatory cytokine interleukin-10 (IL-10) and its receptor IL-10RA. We also analyzed the regulation of IL-10 by flow cytometry on SCARF1^+^ BDCA1^+^ DCs (Fig. 3B) and discovered a significant increase (*p*<0.01) in IL-10 protein expression in these cells. IL10 expression can be impaired if SCARF1 is not present, as shown by the use of a blocking antibody (Fig 3B). In addition, we confirmed our findings of IL-10 and SCARF1 upregulation by qPCR (Fig. 3C-D) and observed a significant increase (*p*=0.02) in the response. These data suggest a role for SCARF1 in the anti-inflammatory response during AC removal by enhancing the migratory effect on phagocytes.

### SCARF1 removal of apoptotic cells regulates the phosphorylation of MAPK and STAT1/2

Our Nanostring data also revealed some differences in the MAP kinase, JAK, and STAT families of kinases (Fig. 4A). Kinases are essential for downstream signaling pathways and dictating the immune response that will be generated, such as IL-10 and IFN*α*. Our data showed upregulation of STAT1 through STAT4 in the presence of ACs. Because phosphorylation of these kinases leads to activation, we used phosphor-flow to assess their activation status (Fig. 4B-I). We observed a significant increase (*p<*0.05) in the phosphorylation in STAT1 Ser727 (Fig. 4B-C), STAT2 Tyr690 (Fig. 4D-E), and MAPKp38 (Fig. 4F-G). STAT1 interacts with STAT3 in the anti-inflammatory response, particularly in the production of IL-10 (32). STAT2 can act alone or as a dimer with STAT1 and is involved in the type I and type III interferon (IFN) responses (33). However, our data does not show the upregulation of the IFN responses (Supplemental Figure 1B). MAPKp38 is activated by stress factors, such as the removal of ACs (34). Furthermore, our data showed no difference in ERK1/2 p44/42 phosphorylation (Fig. 4H-I), suggesting that activation of IL-10 is properly balanced to maintain homeostasis. These findings confirm previous work that showed the regulation of IL-10 in the presence of ACs (35). In addition, previous reports demonstrated that DCs can preferentially secrete IL-10 but not IL-12 through the differential regulation of ERK and p38 pathways to regulate the physiological states of DCs (36, 37). Furthermore, ERK have been shown to negatively regulate STAT1, where activation of ERK leads to the ubiquitination of STAT1 for degradation increasing inflammation (38). Together, these data suggest that SCARF1 plays a role in the activation of kinases that are central to the immune response.

**Figure 4.**
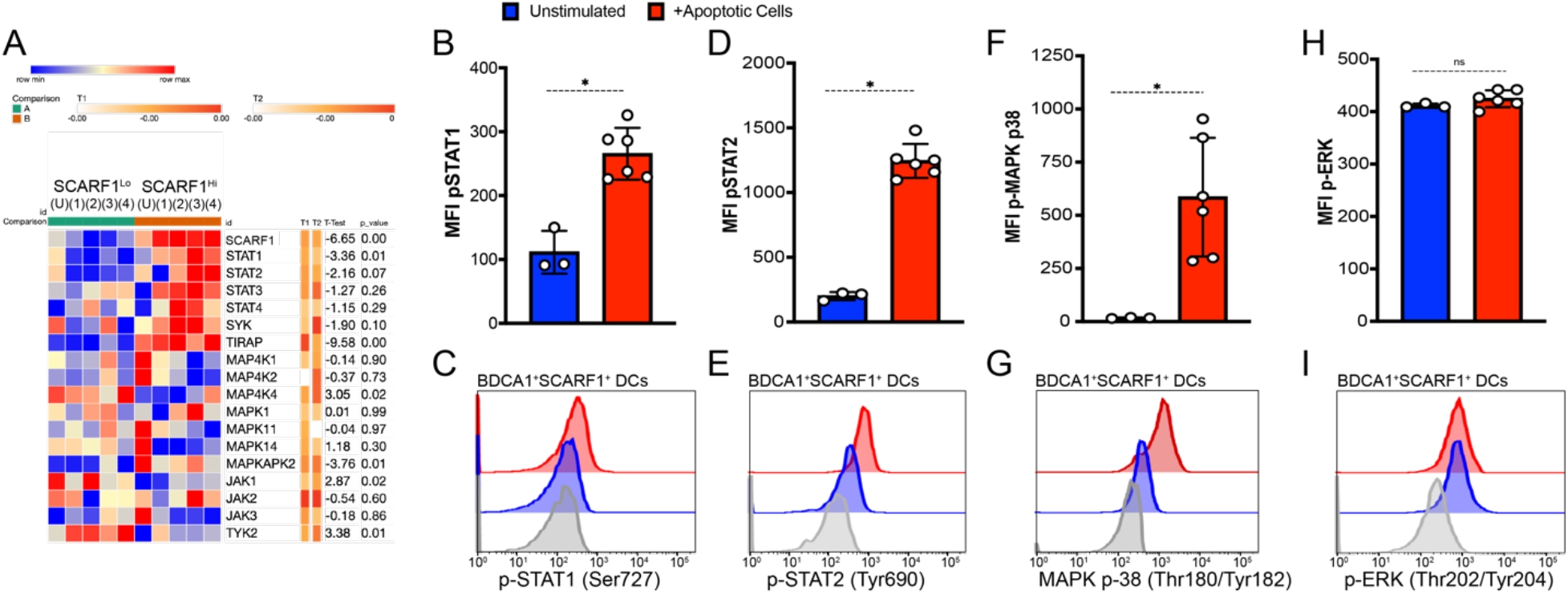
SCARF1 regulates AC-induced phosphorylation of MAPK and STAT. (A) Upregulation of STAT genes in SCARF1^+^ BDCA1^+^ DCs. PBMCs were isolated from healthy controls and treated with apoptotic cells for 4 hours. Live BDCA1^+^ DCs were sorted into SCARF1 negative (SCARF1^Lo^) or SCARF1 positive (SCARF1^Hi^) groups. mRNA was extracted, and the genetic profiling was measure by Nanostring immunology panel. Unbiased heatmap depicting the regulation of kinases. Gene expression as log 2 after normalization using Morpheus software. Representative heatmap of single Nanostring assay (n=4). U, unstimulated. P values by Students *t*-test. (B-I) PBMCs from healthy controls (1×10^6^/mL) were treated with Live/Dead-UV-labeled apoptotic cells for 30 minutes. Cells were immediately fixed and stained for flow cytometry. Cells were gated on CD11c^+^BDCA1^+^SCARF1^+^ in the presence or absence of apoptotic cells. Top panel: MFI quantification, Bottom panel: Representative histograms (B-C) p-STAT1 Ser727, (D-E) p-STAT2 Tyr690, (F-G) p-MAPK p38 Thr180/Tyr182, (H-I) p-ERK p44/42 Thr202/Tyr204. MFI, mean fluorescent intensity. Data represent the mean (±SEM) of 3 independent experiments with 1–2 samples. *P* *(<0.02), *ns* (not significant) by the Mann-Whitney test.

### Lupus patients express similar levels of SCARF1 compared to healthy controls but exhibit altered SCARF1 function and regulation

Our initial observation in our mouse model was that deficiency in SCARF1 results in an increase in ACs, leading to the development of autoimmunity-like lupus disease (13). Furthermore, in Figure 1 we showed that, in healthy controls, SCARF1 is highly expressed on phagocytic cells, specifically DCs and monocytes. We hypothesized that the base levels of SCARF1 would be reduced in lupus patients compared to healthy controls. To answer this question, we obtained SLE patient samples (n=17) and healthy control samples (n=7) and performed flow cytometry to examine the basal levels of SCARF1. We observed increased SCARF1 in plasmacytoid DCs (pDCs) and BDCA1^+^ DCs from a subset of SLE patient samples. This subset of patients had additional phenotypes of lupus vasculitis (n=2) and previous aneurysm (n=1). However, the differences observed were not significant between the SLE patients overall and healthy donors (Fig. 5A). It is important to note that SLE patients were taking medications at the time of the blood draws (Table 1), which could alter protein expression on phagocytes (39, 40).

**Figure 5.**
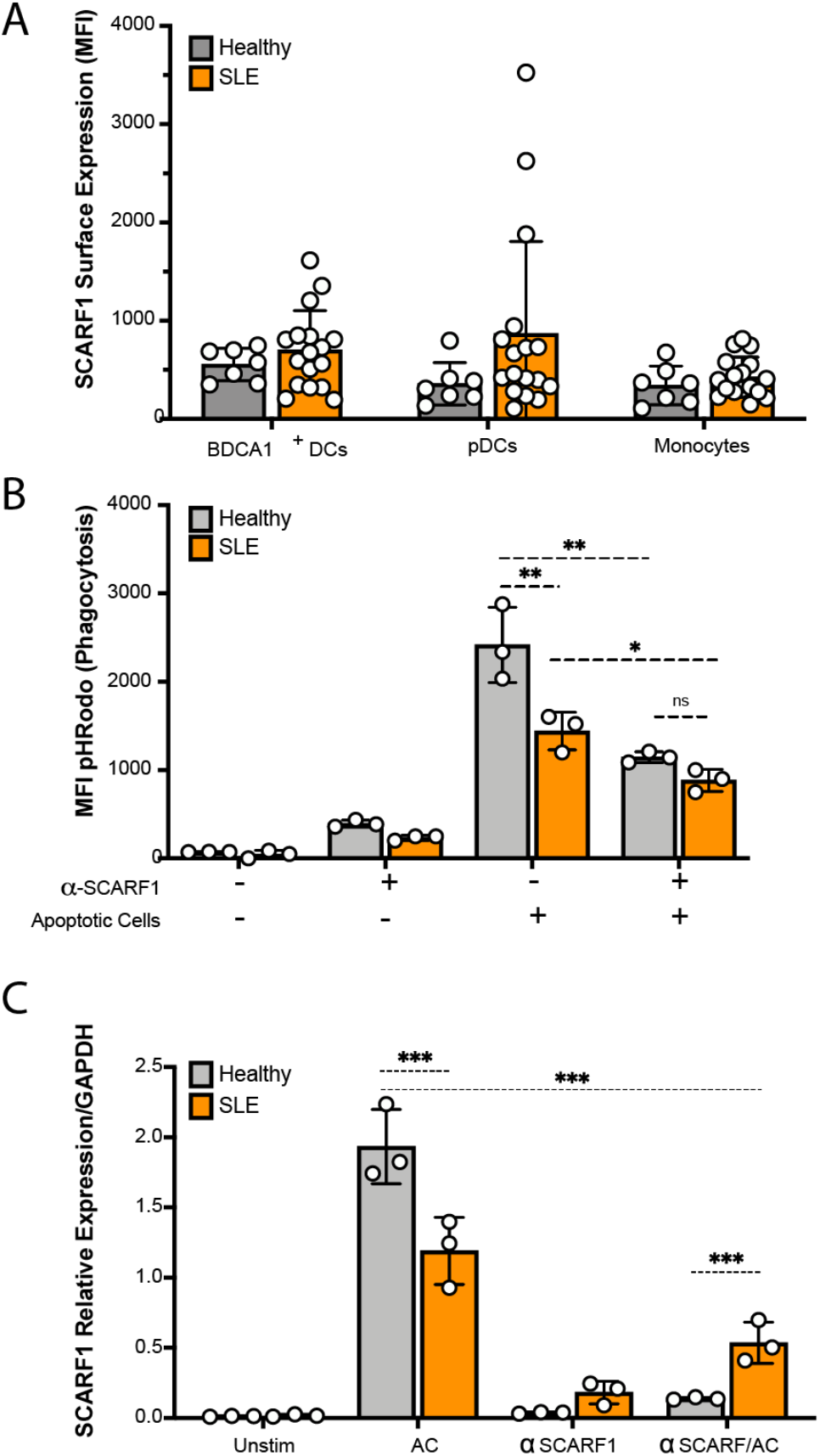
SCARF1 expression is dysregulated after AC uptake. (A) The basal expression of SCARF1 is comparable between SLE patients and healthy controls. PBMCs were isolated from healthy controls (n=7) or SLE patients (n=17). PBMCs were stained for SCARF1, BDCA1^+^ DCs, pDCs, and monocytes. Data are shown as MFI of SCARF1 surface expression. Data are not significant by Mann-Whitney test. (B-C) SCARF1 contributes to the removal of apoptotic cells in Lupus patients. (B) PBMCs (1×10^6^/mL) were incubated with anti-SCARF1 for 30 minutes, then stimulated with pHrodo red-labeled apoptotic cells (2×10^6^/mL). Cells were incubated for 4 hours. Cells were stained and analyzed by flow cytometry for SCARF1 and BDCA1^+^ DCs. Data are the mean (±SEM) of 3 independent experiments with n=1 per group. MFI, mean fluorescent intensity. *P*< *(0.01), **(0.005), *ns* (not significant) by two-way ANOVA with Geissler-Greenhouse correction. (C) BDCA1^+^ sorted DCs (1×10^6^/mL) were incubated with anti-SCARF1 for 30 minutes and then stimulated with apoptotic cells (2×10^6^/mL) for 4 hours. Cells were lysed and mRNA was measured using qPCR. Data are the mean (±SEM) of 3 independent experiments. *** *P*<0.0001 by two-way ANOVA.

**Table 1.** Summary of patient characteristics for subset of SLE patients and healthy controls for PBMCs analyses.

|  | **SLE** |  | **Healthy** |  |
| --- | --- | --- | --- | --- |
| **Sex (n, %)** | Female | Male | Female | Male |
|  | (16, 95%) | (1, 5%) | (7, 100%) | (0, 0%) |
| **Average Age (SD)** | 39 (16) | 23 (0) | 28.8 (5.2) |  |
| **Medication at time of blood draw (n, %)** |  |  |  |  |
| *Glucocorticoids* | 7 (41.2%) |  | UNK |  |
| *Hydroxychloroquine* | 11 (64.7%) |  | UNK |  |
| *Oral immunosuppressant* | 5 (29.4%) |  | UNK |  |
| *Biologic immunosuppressant* | 1 (5.8%) |  | UNK |  |
| **Disease activity (n, %)** |  |  |  |  |
| *Active SLE* | 4 (23.53%) |  |  |  |
| *Remission/Low activity SLE* | 13 (76.47%) |  |  |  |

We wanted to address if vasculitis could be a contributing factor for higher SCARF1 expression. Therefore, we used our Scarf1-deficient mouse model to help answer these questions. We used the *Candida albicans* water-soluble fraction-induced vasculitis model to identify whether SCARF1 plays a role in vasculitis (41). Our data showed no difference between WT and SCARF1-deficient mice in the initiation and development of vascular inflammation (data not shown). These results suggest that vasculitis is not the main cause of the upregulation of SCARF1 in this subset of patients. However, additional studies are needed to confirm these findings, as SLE is a very complex and heterogenous disease, and this was only one model to induce vasculitis.

Due to the similar levels of SCARF1 between samples from healthy and most disease patients, we questioned whether SCARF1 on BDCA1^+^ DCs in lupus patients are capable of removing ACs. We randomly picked PBMCs from SLE patients and analyzed the removal of ACs on BDCA1^+^ DCs by pHrodo red. As previously established (42–44), we observed a significant reduction (*p*<0.01) in the uptake of ACs in lupus patients compared to healthy controls (Fig. 5B). The addition of a SCARF1-blocking antibody to cells from SLE patients resulted in a small but significant reduction (*p*<0.01) of efferocytosis. Similar to the results shown in Figure 2, in healthy controls we see over 50% reduction in the removal of ACs in the absence of SCARF1, demonstrating the importance of SCARF1 in the removal of apoptotic debris. Furthermore, our data show no difference in SCARF1-blocked BDCA1^+^ DCs between healthy controls and lupus patients in the phagocytosis of debris. We also looked at the mRNA regulation of SCARF1 in SLE patients compared to healthy controls (Fig. 5C). These data show that, in the presence of ACs, the upregulation of SCARF1 is significantly reduced (*p*<0.0001) in SLE patients compared to healthy controls. Furthermore, blocking SCARF1 prior to treating BDCA1^+^ DCs with ACs inhibited SCARF1 mRNA upregulation; however, in SLE patients, SCARF1 mRNA is significantly increased (*p*<0.0001) when compared to healthy controls. Our data demonstrate that SLE patients have altered SCARF1 regulation in the presence of ACs, which may contribute to their impairment in AC clearance.

### Lupus patients exhibit significant levels of anti-SCARF1 autoantibodies in the serum

A hallmark of SLE is the presence of autoantibodies to self-antigens. Recently, autoantibodies to the scavenger receptors MARCO and SR-A have been detected in SLE patient serum (45, 46). To investigate the presence of anti-SCARF1, we developed a colorimetric ELISA-based assay against anti-SCARF1 autoantibodies in human serum. Recombinant human SCARF1 protein was generated in-house, and we used commercial anti-human Ig antibodies to detect anti-SCARF1. We discovered for the first time that anti-SCARF1 IgG autoantibodies were detected in 26% (n=38) of SLE patients when compared to healthy controls (Fig. 6A and Table 2). The presence of anti-SCARF1 antibodies was associated with dsDNA antibody positivity (Table 2). These data demonstrate the presence of anti-SCARF1 autoantibodies in SLE patients.

**Figure 6.**
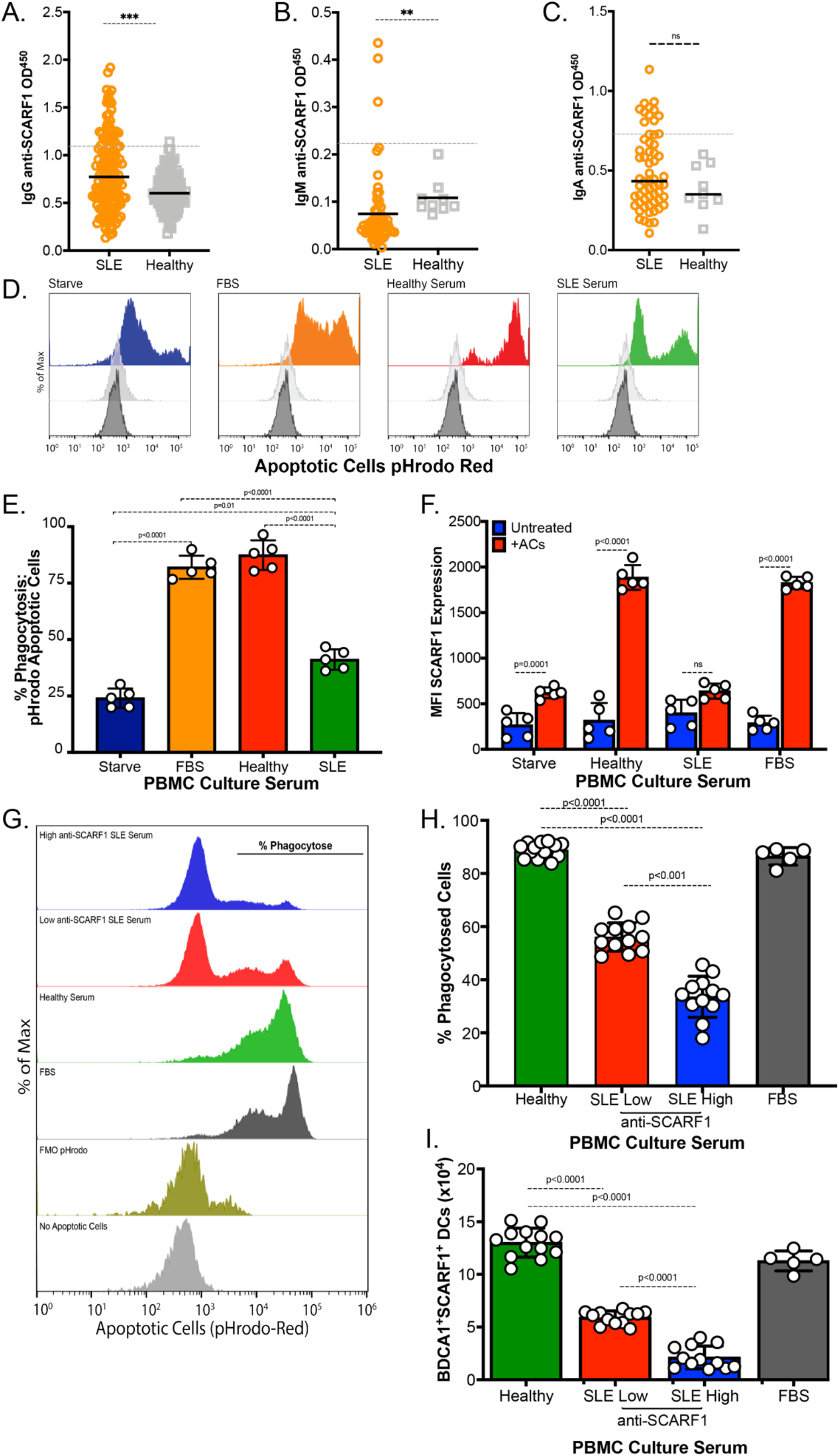
Increased levels of anti-SCARF1 auto-antibodies in the serum of Lupus patients show defects in removal of ACs. (A-C) Soluble SCARF1 or blocking buffer-coated ELISA plates were incubated with sera from SLE patients or healthy individuals. (A) IgG SCARF1 antibodies (Healthy n=100, SLE n=145), (B) IgM SCARF1 antibodies (Healthy n=9, SLE n=62), (C) IgA SCARF1 antibodies (Healthy n=9, SLE n=62). Data are shown as anti-SCARF1 minus anti-block data to reduce the level of binding to the blocking buffer. The optical density (OD) values of each individual are represented as a single point. Dashed lines indicate the OD values that exceed the mean controls by more than 3 standard deviations (sd). Horizontal bars represent the mean values. *P* **(=0.0009), ***(=0.0003), *ns* (not significant) by Mann-Whitney test. (D-F) Lupus serum reduces apoptotic cell uptake in SCARF1^+^ BDCA1^+^ DCs. PBMCs (1×10^6^/mL) were incubated with autologous serum, Lupus serum, or serum starved for 20–24 hours. Cells were incubated for 3 hours with pHrodo red-labeled apoptotic cells (2×10^6^/mL). Cells were stained and analyzed by flow cytometry. (D) Representative histogram of pHrodo expression in SCARF1^+^ BDCA1^+^ DCs. (E) Percentage of ACs phagocytosed by BDCA1+ DCs treated with the specified serum and measured as pHrodo positive cells. (F) MFI quantification SCARF1 expression from BDCA1+ DCs that have phagocytosed ACs. (E-F) Data represent the mean (±SEM) of 3 independent experiments n=5, by Two-way ANOVA. (G-H) Levels of autoantibodies to SCARF1 directly correlates with deficiency in efferocytosis. PBMCs (1×10^6^/mL) were incubated with autologous serum, Lupus serum, or serum starved for 20–24 hours. Cells were incubated for 3 hours with pHrodo red-labeled apoptotic cells (2×10^6^/mL). Cells were stained and analyzed by flow cytometry. (G) Representative histogram of pHrodo expression in SCARF1^+^ BDCA1^+^ DCs. (H) Percentage of pHrodo labeled ACs phagocytosed by BDCA1+ DCs treated with specified serum and measured as pHrodo positive cells. Data represent the mean (±SEM) of 2 independent experiments n=12, by Two-way ANOVA. (I) Autoantibodies block SCARF1 expression on BDCA1^+^ DCs. PBMCs (1×10^6^/mL) were incubated with autologous serum, Lupus serum, or serum starved for 20–24 hours. Cells were incubated for 3 hours with pHrodo red-labeled apoptotic cells (2×10^6^/mL). Cells were stained and analyzed by flow cytometry. Data represent the mean (±SEM) of 2 independent experiments n=12, by Two-way ANOVA.

Next, we investigated the presence of additional anti-SCARF1 autoantibodies, including IgM, which interacts with the classical complement protein C1, and IgA. Anti-SCARF1 IgM autoantibodies were identified in n=3 (2%) SLE patients and no on healthy controls (Fig. 6B), and anti-SCARF1 IgA levels were identified in n=11 (8%) SLE patients and no on healthy controls (Fig. 6C). This data presents novel autoantibodies against SCARF1 in a subset of SLE patients.

### Anti-SCARF1 autoantibodies contribute to reduced efferocytosis in lupus serum

There are additional mechanisms of efferocytosis, including molecules in the serum that can target ACs for removal. Notably, SLE patients suffer from deficiencies in the early complement components such as C1q. C1q and other proteins, including immunoglobulins, influence the removal of apoptotic debris; therefore, such deficiencies can lead to the accumulation of ACs and inflammation (47). Furthermore, we observed the presence of autoantibodies to SCARF1, which we hypothesized may also play a role in abnormal AC clearance. Thus, we hypothesized that treatment with autologous healthy serum would enhance AC uptake compared to serum from SLE patients. However, as previously mentioned, in our assays, C1q did not appear to play an essential role in SCARF1-mediated AC removal. To test this, we used pHrodo red-labeled ACs and assayed for phagocytosis (both uptake and removal) by flow cytometry in serum-free media, healthy serum, and SLE serum containing anti-SCARF1 antibodies (Fig. 6D-E). Our data show that serum-starved BDCA1^+^ DCs and SLE patient serum-treated in BDCA1^+^ DCs exhibit impaired efferocytosis. The serum from these patients show increased anti-SCARF1 IgG and IgM levels. In contrast, healthy serum-treated BDCA1^+^ DCs display effective efferocytosis, indicating a role of anti-SCARF1 in efferocytosis.

We question whether the concentration of autoantibodies to SCARF1 corresponded to the levels of decreased efferocytosis. To test this question, we selected serum from the top 10 highest anti-SCARF1 and the top 10 lowest anti-SCARF1 samples using our ELISA data and compare to healthy controls (Fig. 6G-H). We found that BDCA1+ DCs treated with serum that was expressing high levels of anti-SCARF1 exhibit less efferocytosis, and that this defect correlated with the levels of anti-SCARF1 (Fig. 6G). We discovered up to 50% reduction in removal of ACs when compared to controls (Fig. 6H). Together, these data demonstrate that anti-SCARF1 antibody positive SLE serum inhibits SCARF1 mediated AC uptake, suggesting that autoantibodies to SCARF1 are contributing to blocking the interaction between ACs and SCARF1.

### IgG Depletion increased ACs uptake from lupus serum

Next, we wanted to determine whether autoantibodies to SCARF1 play a role in blocking efferocytosis. In order to determine the role of immunoglobulin in serum, we depleted IgG using Protein A/G agarose beads from healthy and SLE serum and confirm by Western Blot (Supplemental Fig. 3). Next, we assayed for SCARF1 binding using a soluble SCARF1 on Dot Blots (Fig. 7A). Once we demonstrated that IgG were not binding soluble SCARF1, we proceeded to the phagocytosis assay. Depletion of IgG had no effect on healthy controls (Fig. 7B-C; Supplemental Figure 3). However, the data shows a significant increase in efferocytosis after IgG deletion on SLE serum (Fig. 7B-C; Supplemental Figure 3). Our data shows up to a 25% increase in efferocytosis after IgG depletion. Furthermore, we match full and depleted serum samples and looked at the number of BDCA1^+^SCARF1^+^ DCs by flow cytometry. We can observe an increase in phagocytic DCs by up to three-fold numbers of cells (Fig. 7D), suggesting that anti-SCARF1 was blocking SCARF1 protein from this DCs. This data suggests that the serum in SLE contains SCARF1 pathogenic autoantibodies that are blocking or inhibiting access of efferocytosis receptors for the efficient and silent removal of ACs, leading to the accumulation of secondary necrotic cells and inflammation.

**Figure 7.**
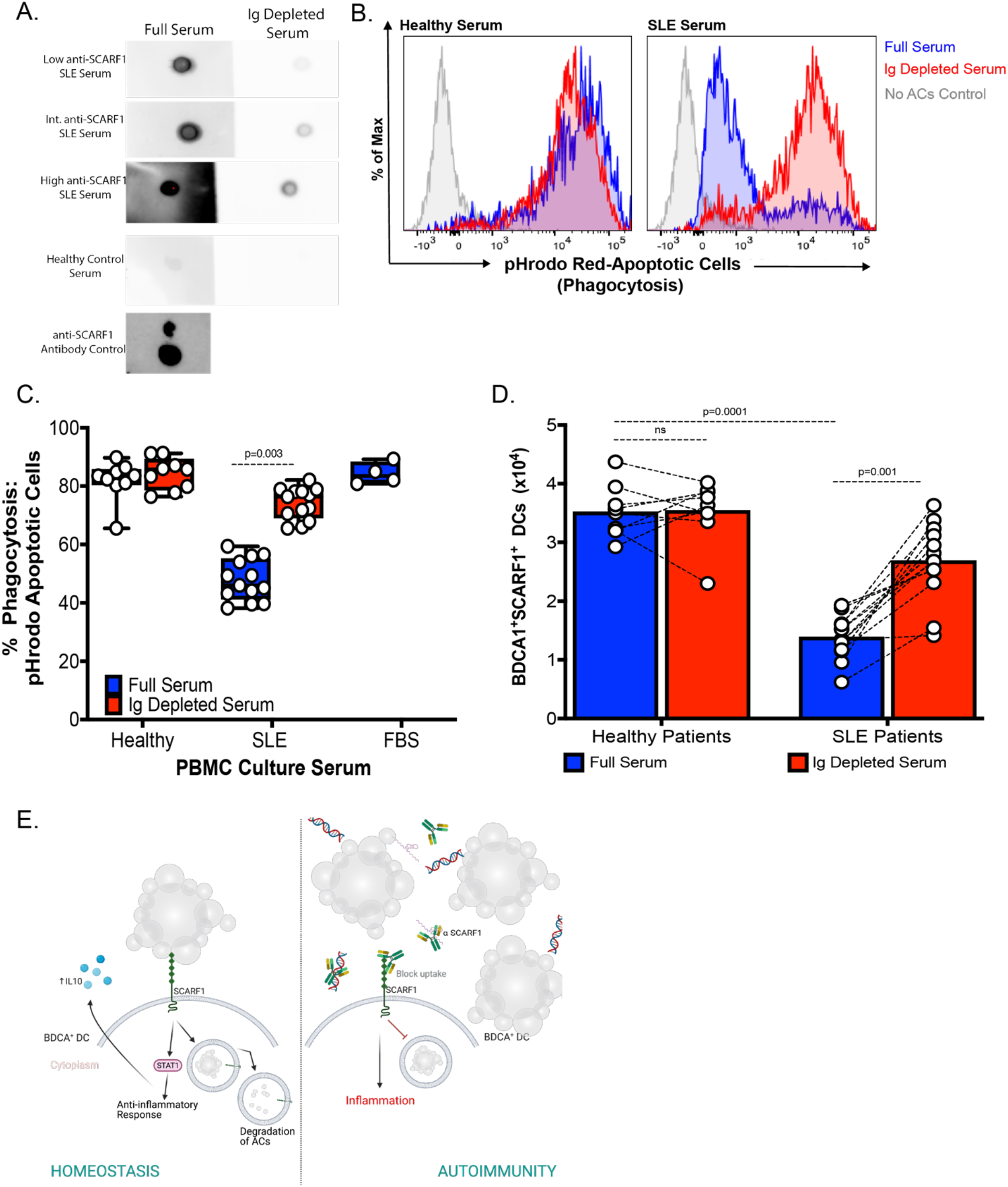
IgG depletion increases SCARF1-mediated ACs clearance. (A) Decreased SCARF1 binding after IgG depletion. We depleted 20% serum in RPMI, healthy or SLE, using 100μL of Protein A/G agarose beads columns. IgG depletion was confirmed by Western Blot (data not shown). SCARF1 binding was analyzed by Dot Blot. Recombinant protein was transfer to nitrocellulose membrane, and full or depleted serum was used as a primary antibody. Human IgG was used as secondary antibody. Representative blot (n=13 SLE and n=8 Healthy). Anti-SCARF1 was use a control. (B-D) Ig-Depletion restores efferocytosis in SLE patient serum. PBMCs (1×10^6^/mL) were incubated with full serum, or Ig-depleted serum for 20–24 hours. FBS was used as serum control. Cells were incubated for 3 hours with pHrodo red-labeled apoptotic cells (2×10^6^/mL). Cells were stained and analyzed by flow cytometry. (B) Representative histograms of pHrodo expression to measure phagocytosis. Blue line, untreated/full serum; Red line, Ig-depleted serum. (C) Quantification of phagocytosed ACs measured as percentage CD11c^+^BDCA1^+^pHrodo^+^ cells. Data represent the mean (±SEM) of 2 independent experiments n=13 SLE and n=8 Healthy, by Two-way ANOVA. (D) Increased BDCA1^+^SCARF1^+^ cells after Ig-Depletion. Total number of CD11c^+^BDCA1^+^SCARF1^+^ measure by flow cytometry in the presence of full serum or Ig-depleted serum. Data represent the mean (±SEM) of 2 independent experiments n=13 SLE and n=8 Healthy, by Two-way ANOVA. (E) Proposed Model of SCARF1 in human SLE development. Diagram made using Biorender.

Our study confirms a role for SCARF1 as an efferocytosis receptor in humans. We propose the following model (Figure 7E): upon interaction of SCARF1 with ACs, the complex is internalized, this results in the production of cytokines and chemokines, essential for the rapid removal of ACs. We demonstrate that IL-10 is produced by the activation and phosphorylation of STAT1 in the SCARF1 pathway. While our model does not show a differential expression of membrane-bound SCARF1 in SLE phagocytes, we did observe a significant increase in the production of anti-SCARF autoantibodies. These antibodies could play a pathogenic role in the removal of ACs by blocking the receptor and inhibiting removal of ACs. We provide evidence suggestive of this mechanism, as the degree of efferocytosis correlated negatively with the level of anti-SCARF1 autoantibodies, and depleting Ig resulted in improved efferocytosis. However, additional studies are needed to elucidate the role of SCARF1 autoantibodies and receptor function, and whether the presence of this autoantibodies can function as biomarkers in SLE.

## Discussion

Immune reactivity to apoptotic debris is tightly controlled to prevent the development of undesirable inflammation (48). We previously reported a role for SCARF1 as a DC receptor for ACs via interactions with C1q/phosphatidylserine complexes on dead cells. Mice deficient in SCARF1 develop lupus-like autoimmune disease, and present symptoms similar to the human disease, such as the spontaneous generation of autoantibodies to chromatin, immune cell activation, dermatitis, and nephritis. This observation raises the question of whether the role of SCARF1 is conserved from mice to humans, and whether SCARF1 plays a role in SLE development (13). We show experimental evidence for the first time in humans that SCARF1, expressed on BDCA^+^ DCs, is a critical receptor in the uptake and removal of ACs. In the present study, we confirm the expression of SCARF1 on monocytes and their role in efferocytosis. Blocking SCARF1 in BDCA1^+^ DCs results in a reduction of AC removal. A similar mechanism has been described where clearance of microparticles formed during apoptosis depends on IFN*α*, and this removal was mediated by SCARF1 expressed on monocytes and monocyte-derived macrophages (49). Thus, we identified a conserved role for SCARF1 in AC removal in both humans and mice and showed that BDCA1^+^ DCs mediate this process.

After antigen recognition, DCs produce an array of cytokines and chemokines necessary to guide the quality and quantity of the immune response (50). We performed genetic profiling of BDCA1^+^ DCs that are SCARF1^HI^ or SCARF1^Lo^. In our model, we observe upregulation of IL-10 and IL-1RN in healthy SCARF1-expressing cells. This process is essential to suppress inflammation and properly regulate the immune response (51). However, IL-10 has been shown to play a dual role in immune responses (52). On one hand, IL-10 is responsible for dampening inflammation and promoting self-tolerance (53). For example, IL-10 play an essential role in the efferocytosis process in macrophages at the engulfment and post-engulfment states (54). On the other hand, IL-10 was shown to play a pathogenic role in SLE by acting as a growth factor on cytotoxic lymphocytes and promoting survival of autoreactive B cells (52). A recent study reported a reduction in IL-10 on SCARF1-deficient mice during *Mycobacterium tuberculosis* infection (55). *M. tuberculosis* induces apoptosis and/or necrosis dictating the outcome of the infection (56), suggesting a role for SCARF1 in regulating pulmonary cessation and necrosis, which are associated with poor treatment outcomes (55). Our study identifies a new role for SCARF1 in autoimmune disease pathogenesis.

Previous studies suggest that members of the STAT family play a role in the pathogenesis of SLE (57, 58). The *STAT1-STAT4* locus has been associated with increased lupus disease risk (59). During AC removal, we observed the upregulation and phosphorylation of STAT1 and p38 MAPK. However, we did not observe phosphorylation of p42/44 ERK. This finding confirms work by Chung *et al*., where they showed that IL-10 production stimulated by ACs is regulated by p38 MAPK and partially mediated by CD36 by cell-cell contact, but independent of phagocytosis (35). STAT signaling is known to be essential for the activation of IL-10 and type I IFN. Gain of function mutations increase the type I and type III IFN responses, however, at basal levels STAT1 collaborates with STAT3 for the immunomodulatory response-including the activation of IL-10 (60). Furthermore, in macrophages, STAT1 and STAT3 have been shown to polarized M1 and M2 during efferocytosis by inducing IL4, IL-10 and low levels of IFN*γ* for M2 (61). Therefore, STATs are responsible for the fine tuning between inflammation and the anti-inflammatory response. Xu *et al*. recently showed on a liver transplant model using rats, that SCARF1 promotes M2 polarization of Kupffer cells by enhancing phagocytosis via the PI3K-AKT-STAT3 signaling pathway (62). In our model, phagocytosis of ACs by SCARF1 was required for IL-10 production via STAT1 phosphorylation. Thus, SCARF1 not only acts as a receptor for ACs but is also required for the production of a key anti-inflammatory cytokine.

Our initial observations using a mouse model was that deficiency in SCARF1 results in the development of autoimmune disease. This led us to hypothesize that SCARF1 is dysregulated in SLE patients. We did not observe a significant difference in SCARF1 expression on phagocytic cells, including BDCA1^+^ DCs for overall SLE patients versus healthy controls. However, we did observe the novel identification of autoantibodies to SCARF1. Autoantibodies are a known hallmark of SLE, and they can precede the onset of autoimmune disease symptoms (63). Consistent with this observation, the presence of autoantibodies to scavenger receptors have been shown in lupus-prone mouse models (64). Chen *et al*. discovered the presence of anti-MARCO and anti-scavenger receptor A in 18.5% of SLE patients (45). Recently, Zhou *et al*. showed that anti-Tyro3 was associated with disease activity and the presence and impaired efferocytosis (65). We observe IgG (26%), IgM (2%) and IgA (11%) anti-SCARF1 antibodies in of SLE patients. Furthermore, we also observed a reduction in the phagocytosis of ACs that were treated with SLE serum on SCARF1^+^ BDCA1^+^ DCs. This novel finding was demonstrated in Figure 6D-E by the addition of exogenous anti-SCARF1, where we observed over 50% reduction in efferocytosis. This defect in efferocytosis correlated with the levels of anti-SCARF1 expressed on BDCA1^+^ DCs implying that autoantibodies are blocking SCARF1 and are therefore pathogenic. Growing evidence suggests that serum autoantibodies may play a role in the impairment of AC removal by showing reactivity to molecules on the AC and/or the phagocytes (45, 66). Similar to our findings, multiple groups have observed impaired efferocytosis on SLE patients and was primarily due to the presence of IgG antibodies (67, 68). Further studies are needed to demonstrate that anti-SCARF1 autoantibodies are responsible for the defect in efferocytosis in SLE patients. However, our data demonstrates that autoantibodies are responsible for phagocyte dysfunction by blocking SCARF1 on BDCA1^+^ DCs (Fig. 7D; Supplemental Figure 3). Altogether, the results indicate that autoantibodies to SCARF1 might be involved in the breakdown of self-tolerance by impairing the uptake of ACs, and, therefore, play a role in the development of SLE.

One limitation of our study is that SLE patients were taking medications at the time of sample collection (Table 1 and Supplemental Table 1), potentially altering the levels of SCARF1 present in cells. Medications (e.g., hydroxychloroquine), steroids (e.g., prednisone), and disease-modifying antirheumatic drugs (e.g., methotrexate) are immunosuppressive drugs that can interfere with DNA synthesis and prevent cells in the immune system from dividing. Alternately, some of the medication the patients were taking could upregulate SCARF1. Hydroxychloroquine is an alkalinizing lysosomotropic drug that accumulates in lysosomes and inhibits important functions by increasing the pH (69). We hypothesize that a drug like hydroxychloroquine will result in the accumulation of membrane-bound SCARF1 on the cell surface of phagocytes by inhibiting the phagocytosis process. Therefore, further studies are needed to determine the impact of drugs on SCARF1 regulation to determine if a reduction in SCARF1 is involved in the pathogenesis of SLE.

In conclusion, this study demonstrates for the first times in humans that SCARF1 plays an important role in efferocytosis and in the development of lupus by modulating the efferocytosis process. We demonstrated that the function of SCARF1 is conserved from worms to mouse to humans as an efferocytosis receptor (13, 70). Not only does SCARF1 efficiently recognize and capture AC, but SCARF1 is responsible for initiating an anti-inflammatory response. A proposed mechanism for altered SCARF-1 mediated efferocytosis in lupus is the presence of autoantibodies to SCARF1 in a subset of SLE patients, which correlate with the presence of anti-dsDNA autoantibodies. The defective clearance of AC in SLE patients appears to depend on the occurrence of such autoantibodies, where SCARF1 was not accessible on BDCA1^+^ DCs to uptake the apoptotic cells. Such failure to remove apoptotic debris will lead to the accumulation of immunogenic noxious molecules leading to a vicious cycle of inflammation and autoimmunity. Understanding the mechanisms of apoptotic cell clearance in healthy and disease states can shed light in novel therapeutic strategies.

## Materials and Methods

### Reagents and Cell Culture

Reagents were purchased from Sigma-Aldrich unless otherwise stated. Complete media consisted of RPMI-1640 (Gibco, Thermo-Fisher, Grand Island, NY) supplemented with 100 U/mL penicillin, 100 U/mL streptomycin, 2 mM Glutamax and 10% fetal bovine serum (FBS, Hyclone GE Lifescience, Pittsburgh, PA). TIME cells (Cat#: CRL-4025) were obtained from the ATCC (Manassas, VA). Culture of the cell line required Vascular Cell Basal Medium (ATCC, Cat#: PCS-100-030), supplemented with microvascular endothelial cell growth kit-VEGF (ATCC, Cat#: PCS-110-041). Mouse embryonic fibroblasts (MEFs; Cat# SCRC-1008) were obtained from the ATCC. Flow cytometry antibodies were obtained from BD Biosciences. Human anti-SCARF1 (Cat#: AF2409 and MAF2409) antibodies were purchased from R&D Systems.

### Human PBMC isolation and patient samples

Peripheral blood was obtained from healthy adult donors under an Institutional Review Board (IRB)-approved procedure by the Massachusetts General Hospital (MGH) Blood Bank. Peripheral blood from SLE patients was obtained from the Rheumatology Practice at MGH, with IRB approval. In addition, we obtained serum from 145 patients with SLE, including 20 from a rheumatology practice at Massachusetts General Hospital and 125 from the large, multicenter Partners Healthcare Biobank. SLE diagnoses were confirmed on expert MD review, and all subjects met the 1997 American College of Rheumatology classification criteria for SLE. We obtained serum from 100 healthy controls from the Biobank.

Human PBMCs were isolated as described (71, 72). Briefly, a portion was clotted, and the autologous serum was collected following centrifugation. The remainder of the blood was anticoagulated with heparin, and the PBMCs were purified by Ficoll-Hypaque (GE Healthcare) density gradient centrifugation for 30 minutes at room temperature, 400x*g* no brake. Cells were collected and treated as specified. For some experiments, human BDCA1-dendritic cells were positively selected using CD1c^+^ (BDCA1; Miltenyi Cat#: 130-119-475) magnetic beads prior to stimulation.

### Nucleofection and CRISPR-*Cas9*

Endothelial cells were transfected using the Nucleofector primary mammalian endothelial cell kit (Cat. #VAPI-1001, Lonza Biosciences, Durham, CA), according to manufacturer’s instructions. Dual CRISPR-*Cas9* (pCLIP-sgRNA and pCLIP-*Cas9*) targeting SCARF1 or control was purchased from Transomics Technologies (Transomic Technologies Inc., Huntsville, AL).

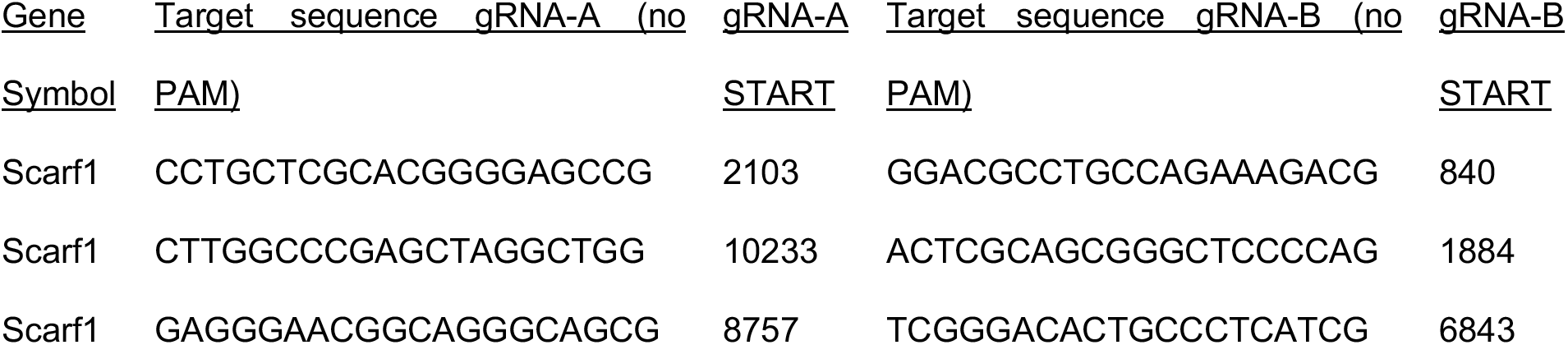

For pCLIP-sgRNA and pCLIP-*Cas9* preparation, bacterial cultures from the stock were propagated in LB media supplemented with 100 μg/mL of carbenicillin until the culture appeared to be turbid. Plasmids were extracted using anendotoxin-free kit (QIAgen Cat. #12362) and stored until ready to use.

Endothelial (TIME) cells were grown to 85% confluency, and 2 x 10^6^ cells were co-transfected with pCLIP-sgRNA and pCLIP-*Cas9* using a Nucleofector II with the program M-003. As a transfection control, we nucleofected GFP alone (p-MAX-GFP, Amaxa). Viable SCARF1-deficient cells were sorted 24hr post-nucleofection by pCLIP-sgRNA-GFP and p-CLIP-*Cas9*-RFP. Experiments were performed at 24hr post-sorting and analyzed by flow cytometry.

### Generation of Apoptotic Cells

Apoptotic cells from MEFs or Jurkat cells were generated following osmotic shock as previously described (13). For all assays, unless specified, apoptotic cells were labeled with live/dead fixable stain or pHrodo. Briefly, cells were first harvested and centrifuged at 800x*g* for 5 minutes. Cells were washed with PBS, then resuspended in hypertonic media (DMEM supplemented with 10% w/v polyethylene glycol 1000, 0.5M Sucrose, 10 mM Hepes) for 10 min at 37 °C/5% CO_2_. Then 30 mL of hypotonic media (60% DMEM, 40% water) was added and incubated for 5 min at 37 °C/5%CO_2_. Cells were washed and incubated with DMEM-C for 4 hours. Apoptotic cells were confirmed using flow cytometry by Annexin/PI stain.

### Phagocytosis of Apoptotic Cells

To measure the phagocytosis of apoptotic cells, we used pHrodo red (Molecular Probes, ThermoFisher; Cat. #P35372). Apoptotic cells were labeled with pHrodo red according to the manufacturer’s instructions (13). Briefly, 5×10^6^ apoptotic cells were collected and washed twice with PBS. Cells are resuspended in 1 mL of PBS with 1 µL of pHrodo stock and incubated for 30 minutes at RT in the dark. pHrodo-labeled cells are washed three times with PBS and resuspended in RPMI media. Labeled apoptotic cells were added at a ratio of 1:1 to PBMCs and incubated up to 4 hrs. Phagocytosis was measured by the increase in fluorescence using a BD LSRII Fortessa and data analyzed using FlowJo 10.6 software version for Mac.

### Microscopy

Cell imaging was performed by confocal microscopy. Cells were cultured in 48-well plates until they reached 70% confluency. Next, the Red Live/Dead (Alexa546) labeled-apoptotic cells were added to the wells for 2 hrs. The wells were carefully washed with fresh media, and warm RPMI-C media was added. Cells were incubated for a total of 2 hrs, and then fixed, washed, and immobilized to a microscope slide by cytospin. Intracellular staining was performed for SCARF1. Cells were stained with DAPI. Chambers were removed and fixed with coverslips until ready to analyze. Slides were visualized using a Zeiss LSM Confocal Microscope, and data were analyzed using Zeiss Zen Blue Software (Version 3.1).

### Real-time quantitative PCR

Total RNA was extracted using the RNeasy kit according to the manufacturer’s instructions (QIAgen). Each sample was reverse transcribed using multiscribe reverse transcriptase (Applied Biosystems, Foster City, CA). Each qPCR reaction was 25 μL and contained 2 μL of cDNA, 12.5 μL of 2x SYBR green master mix (Applied Biosystems), 500 nM of sense and antisense primers, and PCR-grade water. Oligonucleotide primer sequences were designed on the PrimerBank website from Integrated DNA Technologies (IDT, Coralville, IA). The following primers were used:

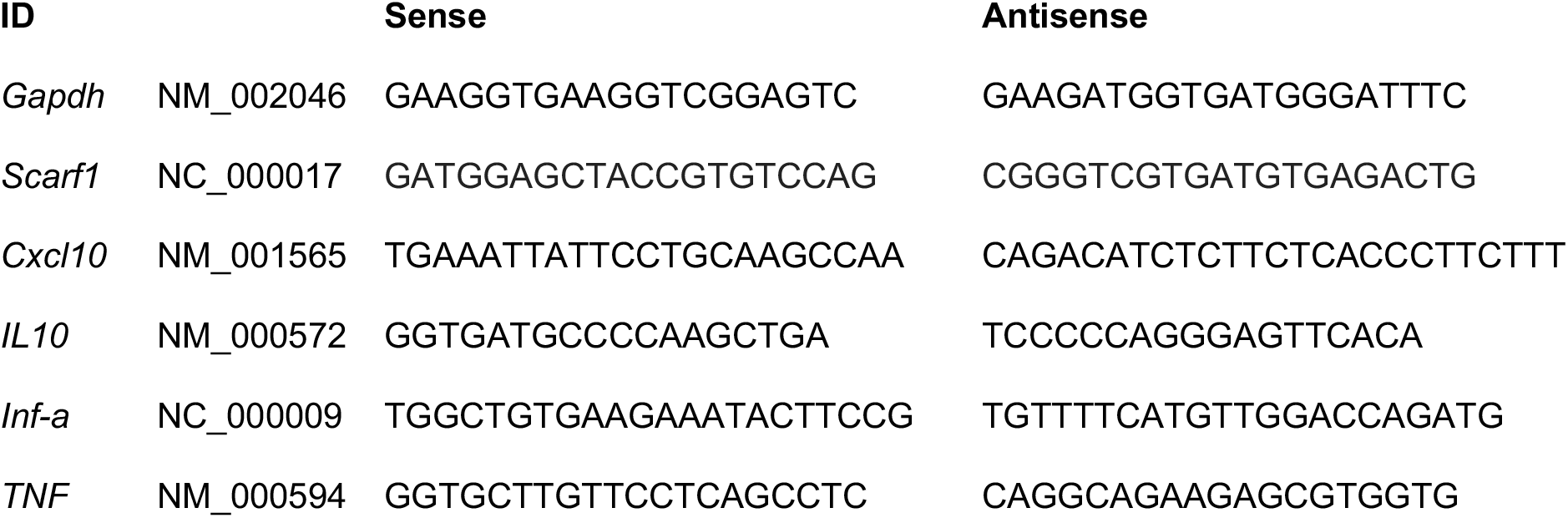

For each individual sample, emitted fluorescence was measured three times during the annealing-extension phase using the LightCycler 96 System (Roche, Basel). Amplification plots were analyzed using the LightCycler 96 System Application Software (Roche). Gene expression was quantified by comparing the fluorescence generated by each sample with a standard curve of known quantities. The calculated number of each sample was divided by the number of housekeeping gene *Gapdh*.

### nCounter Analysis

Nanostring assay was performed according to the manufacturer’s protocol (NanoString Technologies). Probe sequences for human immunology were designed and manufactured by NanoString. Each CodeSet included a number of housekeeping genes selected from a publicly available database to ensure RNA quality differences. Briefly, cells were treated and sorted as specified above. RNA was extracted using RNeasy Plus Micro Kit (QIAgen Cat. # 74034), and the concentration of RNA was measured by Qubit Fluorometric Quantification (ThermoFisher). The master mix and the hybridization reaction for the code sets were generated according to the manufacturer’s protocol. The next day, hybridized reactions were loaded on the Sprint cartridges. Samples were analyzed using the nCounter SPRINT Profiler. Generated data went through an internal QC process using the housekeeping genes, and secondary analysis included normalization of the genes using nSolver Data Analysis. Heatmap was generated using Morpheus Software adjusted for the Log2 and Z-score values (Broad Institute, https://software.broadinstitute.org/morpheus/).

### Flow Cytometry

PBMCs were isolated as described above. To properly identify SCARF1-expressing cells, we designed a multicolor cytometry assay, described below.

| **Antibody** | **Fluorophore** | **Clones** | **Cell Marker** | **Company** |
| --- | --- | --- | --- | --- |
| BDCA2 | BV421 | V24-785 | DCs, conventional 2 | BD Biosciences |
| CD123 | BV510 | 9F5 | DCs, plasmacytoid | BD Biosciences |
| CD14 | Alexa700 | M5E2 | Monocytes (CD14<sup>+</sup>CD16<sup>-</sup>) | BD Biosciences |
| CD16 | PE-Cy7 | 3G8 | Monocytes | BD Biosciences |
| CD11c | PerCP-Cy5.5 | B-ly6 | All dendritic cells | BD Biosciences |
| CD8 | BUV737 | SK1 | CD8<sup>+</sup> T cells | BD Biosciences |
| CD4 | BUV395 | SK3 | CD4<sup>+</sup> T cells | BD Biosciences |
| CD3 | BV786 | HIT3α | T cells | BD Biosciences |
| CD66 | BV605 | B1.1/CD66 | Neutrophil | BD Biosciences |
| BDCA1 | PE | F10/21A3 | DCs, conventional 1 | BD Biosciences |
| BDCA3 | FITC | 1A4 | DCs, plasmacytoid | BD Biosciences |
| CD11b | BV711 | ICRF44 | DCs and monocytes | BD Biosciences |
| SCARF1 | Alexa647 |  | All cells | R&D Systems |
| CD45 | PE CF594 | 2D1 | All lymphocytes | BD Biosciences |
| CD19 | APC Cy7 | SJ25C1 | B cells | BD Biosciences |

Single color controls and fluorescence minus one (FMO) were used as controls and to set up the proper voltages. Samples were collected using a 5-laser BD Fortessa (BD Biosciences) and BD Diva Software. Data were analyzed using FlowJo X version for Mac.

### Phospho-flow cytometry analysis

PBMCs were isolated as described above. BDCA1^+^ DCs were sorted using magnetic sorting using the BDCA^+^ kit from Miltenyi (Miltenyi Biotech, Cat. #130-119-475) and allowed to recover overnight. BDCA^+^ DCs (2×10^6^ cells/mL) were seeded in 5 mL polypropylene tubes containing RPMI complete media. Cells were treated with a 1:1 ratio of apoptotic cells to DCs or left untreated and incubated 25 minutes at 37 °C with 5% CO_2_. Treated cells were fixed with cold 2% paraformaldehyde for 10 minutes at 4 °C. Cells were washed three times with cold FACS buffer (PBS supplemented with 2% FCS) and permeabilized with cold 100% methanol for 20 minutes on ice. Cells were washed twice to remove methanol and stained with fluorescently labeled antibodies, Live/dead stain-UV (Molecular Probes), SCARF1-APC (R&D Systems, Cat. #FAB2409R), CD11c-, BDCA1^+^-PE (Miltenyi), pSTAT2-Alexa488 (Tyr690, Cell Signaling clone D3P2P), p-STAT1-Alexa488 (Tyr701, Cell Signaling clone 58D6), Erk1/ 2 p-44/42-Alexa488 (Cell Signaling clone 137F5), and MAPK p38-Alexa488 (Thr180/Tyr182, Cell Signaling clone 3D7). Samples were analyzed using BD LSRII-Fortessa, and data collected using BD FACSDiva Software and analyzed using FlowJo X version for Mac.

### Recombinant Scarf1 protein expression and purification

*E. coli* (BL21) cells were transformed with an expression plasmid containing Scarf1 fused in frame with a His-tag. Selected clones were grown at 30 °C in LB medium containing 100 μg/mL of ampicillin, and the expression of Scarf1 protein was induced by isopropyl-D-thio-galactoside (IPTG). The bacteria were harvested by centrifugation at 6,000 rpm, and the pellet was resuspended in lysis buffer with protease inhibitor cocktail (Roche, Germany). Then, the cell suspension was lysed by freeze-thawing and incubated with lysozyme (100 mg/mL) and RNase (5 g/mL). Next, the lysate was sonicated ten times with 10 s intervals. Scarf1 protein was purified by affinity chromatography using Ni-NTA columns followed by imidazole elution and confirmed by Western blot (Supplemental Figure 2A).

### Anti-SCARF1 ELISA in patient serum

We performed a novel sandwich ELISA to identify the levels of anti-SCARF1 in the serum. We coated 96-well plates with 250 ng/well of recombinant SCARF1 made in-house (see above) overnight. The next day, the plate was washed and blocked with 2%BSA in BBT (Borate Buffer Saline pH 8.2, SIGMA Cat# 08059-100TAB-F, 8 mL Tween 20) for 1 hr/RT. Patient sample sera were diluted 1:200 in BBT and FBS. Diluted samples were incubated for 1 hr at RT. Plates were washed five times, and anti-human IgG secondary antibody was added at a 1:10,000 dilution and incubated for 30 min at RT. The plate was washed 5x and the developer reagent was activated by adding 50 μL/well of TMB substrate and 50 μL/well of 0.2M sulfuric acid to stop the reaction. Plates were read on a Spectrophotometer ELISA plate reader. To classify positive value, we used a cut-off value defined as the control median plus three standard deviations.

### Serum IgG Depletion

We use Protein A/G Plus (Pierce, ThermoFisher) Agarose beads, according to manufacturer’s instructions. Briefly, in a microcentrifuge tube combine 100μL of bead slurry with 500μL of media to equilibrate the reaction. Centrifuge and remove the supernatant. Add 20% sample serum in 500μL of media, incubate the reaction at 4 degrees in a shaker for 20 minutes. Centrifuge and transfer the supernatant to a new tube with equilibrated slurry, repeat the process three times. Save the Ig depleted supernatant. SDS-PAGE and Western Blot was used to confirm Ig depletion. To assay for SCARF1 binding, recombinant SCARF1 was run on SDS-PAGE and transfer to a nitrocellulose membrane. Full and depleted serum were used as primary antibody and human IgG (Millipore) was used as secondary antibody. Anti-SCARF1 (R&D Systems, Minneapolis MN) was used as a positive control. Samples were develop using Super Signal West Pico Chemiluminescent (Thermo Scientific, Waltham MA) and visualized using ChemiDoc MP Imaging System (BioRad, Hercules CA). Once confirmed that the autoantibodies to SCARF1 were removed, we use the full serum and Ig depleted serum to analyzed for efferocytosis as described above (see *Phagocytosis of apoptotic cells*).

### Statistics

Statistical calculations were performed using the statistical software GraphPad Prism 8.4. For comparison of two groups, means ±SE were analyzed by two-tail unpaired Students *t*-test with Bonferroni corrections applied when making multiple comparisons. Patient data were analyzed by Student’s *t*-test and *Chi*-square test. Working model created with BioRender.com

### Study Approval

The study was approved by the Massachusetts General Hospital and Partner’s HealthCare System IRB (Protocol #: 2006P002256). Informed consent forms were collected at the time of blood draw following the Partners Human Research Committee Policy and Guidance.

## Author Contributions

Z.G.R.O. designed and performed experiments, analyzed data, and wrote the manuscript. A.M.J. designed and performed experiments, analyzed data, and edited the manuscript. T.L., R.K., S.L. and T.K.M. developed required models and/or performed experiments and/or edited the manuscript. A.M.J. consented and collected human subjects. J.E.K. and A.D.L. provided advice on experimental design and data analysis.

## Acknowledgments

We would like to thank Dr. Melanie Trombly, Assistant Professor at University of Massachusetts Medical School, for help with editing the manuscript. We also would like to thank the Clinical Immunology Laboratories and the Rheumatology Clinic at Massachusetts General Hospital for providing deidentified SLE samples. Dr. Chie Miyabe for performing a vasculitis mouse model (data not shown).

