## Supplemental Table and Figures for "SCARF1-induced efferocytosis plays an immunomodulatory role in humans, and autoantibodies targeting SCARF1 are produced in patients with systemic lupus erythematosus"

**Supplementary Table 1. Prevalence of SCARF1 in patients with systemic lupus erythematosus**

| | Overall | Scarf1 ab positive | Scarf1 ab negative | p value |
|---|---|---|---|---|
| | 146 | 30 | 116 | |
| **Female, n (%)** | 131 (90) | 30 (100) | 101 (87) | 0.04 |
| **Age, mean (SD)** | 45 (14.5) | 42 (16.5) | 45 (13.9) | 0.34 |
| **SLE features, n (%)** | | | | |
| *Rashes/ alopecia* | 87 (60) | 17 (57) | 70 (60) | 0.41 |
| *Photosensitivity* | 32 (22) | 5 (17) | 27 (23) | 0.45 |
| *Arthritis* | 107 (73) | 20 (67) | 87 (75) | 0.36 |
| *Hematologic* | 51 (35) | 13 (43) | 38 (33) | 0.28 |
| *Serositis* | 40 (27) | 9 (30) | 31 (27) | 0.72 |
| *Oral ulcers* | 22 (15) | 4 (13) | 18 (16) | 0.77 |
| *Nephritis* | 46 (32) | 10 (33) | 36 (31) | 0.81 |
| *Antiphospholipid antibody syndrome* | 23 (16) | 4 (13) | 19 (16) | 0.68 |
| *Cardiovascular* | 9 (6) | 2 (7) | 7 (6) | 0.90 |
| *Neurologic* | 16 (11) | 2 (7) | 14 (12) | 0.40 |
| *Raynaud's* | 39 (27) | 6 (20) | 33 (28) | 0.35 |
| *Vasculitis* | 6 (4) | 0 | 6 (5) | 0.35 |
| **Serology (n, %)** | | | | |
| *ANA* | 138 (95)* | 29 (97) | 109 (94) | 0.56 |

| | | | | |
|---|---|---|---|---|
| *dsDNA* | 91 (62) | 25 (83) | 66 (57) | <0.01 |
| *Smith* | 57 (39) | 19 (63) | 38 (33) | <0.01 |
| *RNP* | 68 (47) | 20 (67) | 48 (41) | 0.01 |
| *SSA (Ro)* | 46 (32) | 11 (37) | 35 (30) | 0.49 |
| *SSB (La)* | 24 (16) | 5 (17) | 19 (16) | 0.97 |
| *Antiphospholipid antibodies* | 37 (25) | 4 (13) | 33 (28) | 0.09 |
| *Hypocomplementemia* | 65 (45) | 15 (50) | 50 (43) | 0.50 |
| **Current SLE activity, n (%)** | | | | |
| *Active* | 45 (31) | 9 (30) | 36 (31) | 0.91 |
| *Remission or Low disease activity* | 78 (53) | 17 (57) | 61 (53) | 0.69 |
| *Unknown* | 23 (16) | 4 (13) | 19 (16) | 0.68 |
| **Current SLE Medications, n (%)** | | | | |
| *Glucocorticoids* | 45 (31) | 12 (40) | 33 (28) | 0.22 |
| *Hydroxychloroquine* | 83 (57) | 20 (67) | 63 (54) | 0.22 |
| *Oral immunosuppressant* | 50 (34) | 7 (23) | 37 (32) | 0.36 |
| *Biologic immunosuppressant* | 13 (9) | 3 (11) | 10 (9) | 0.81 |
| ***missing value in 7*** | | | | |

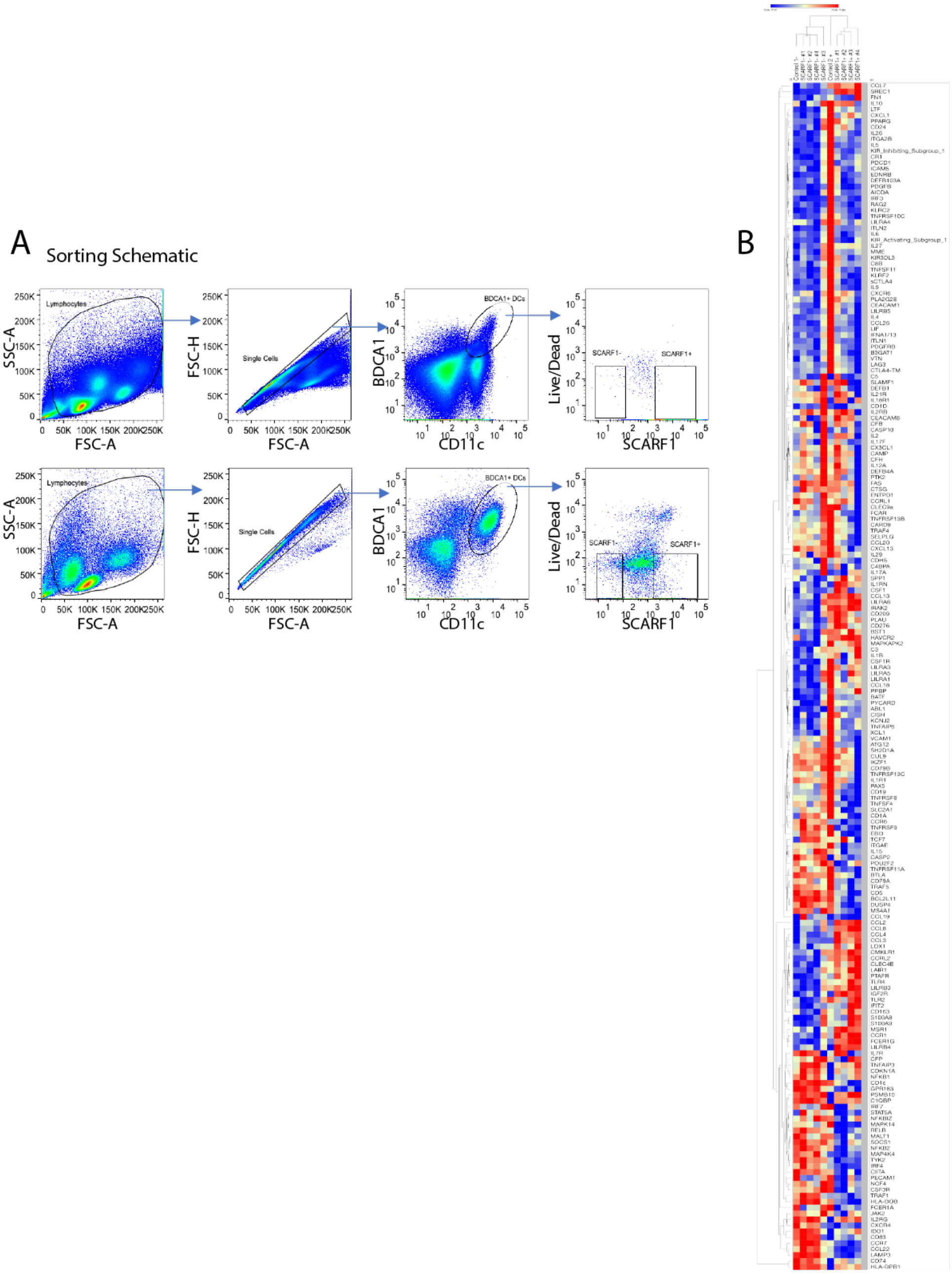


**Supplemental Fig. 1. Differentially expressed genes in SCARF1$^{Lo}$ and SCARF1$^{Hi}$ cells.** (A) PBMCs (2x10$^{6}$) were incubated with apoptotic cells (2x10$^{6}$) for 4 hours. Cells were stained for flow cytometry and sorted into RNA lysis buffer. Shown is a representative experiment out of n=4. (B) Heatmap of Top 150 Nanostring genes. Live BDCA1$^{+}$ DCs were sorted into SCARF1 negative (SCARF1$^{Lo}$) or SCARF1 positive (SCARF1$^{Hi}$) cells. mRNA was extracted, and the genetic profiling was measured using a Nanostring immunology panel. Gene expression as log 2 after normalization using Morpheus software. Shown is a representative heatmap of a single Nanostring assay (n=4). U, unstimulated. P values by Student's *t*-test.

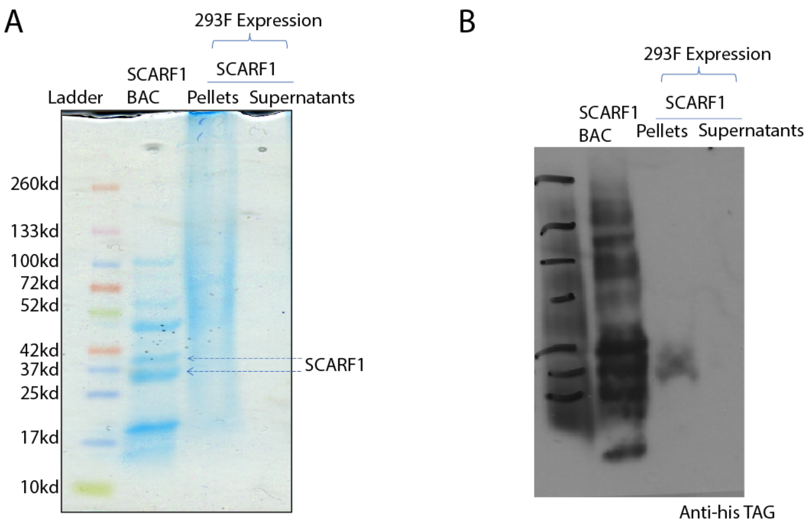


**Supplemental Fig. 2. Expression of Soluble SCARF1 for the auto-antibody ELISA assay.**
(A) SDS Page gel of SCARF1 expression (B) Western Blot of SCARF1 expression.

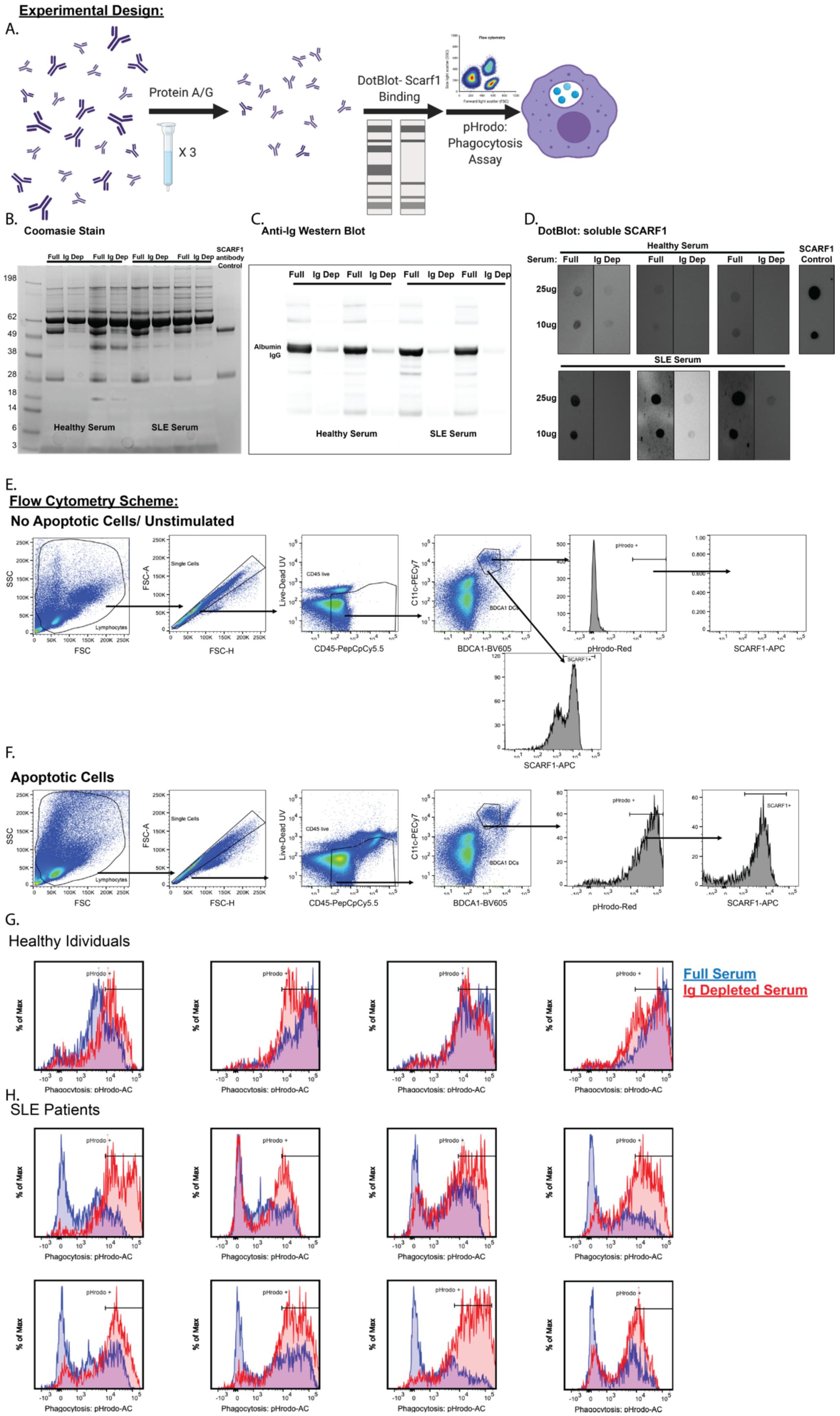
Experimental Design:
A.
Protein A/G
X 3
DotBlot- Scarf1 Binding
Flow cytometry
pHrodo: Phagocytosis Assay
B.
Coomasie Stain
Full Ig Dep Full Ig Dep Full Ig Dep Full Ig Dep
SCARF1 antibody Control
198
62
49
38
28
18
14
6
3
Healthy Serum
SLE Serum
C.
Anti-Ig Western Blot
Full Ig Dep Full Ig Dep Full Ig Dep Full Ig Dep
Albumin
IgG
Healthy Serum
SLE Serum
D.
DotBlot: soluble SCARF1
Healthy Serum
Serum: Full Ig Dep Full Ig Dep Full Ig Dep
SCARF1 Control
25ug
10ug
SLE Serum
25ug
10ug
E.
Flow Cytometry Scheme:
No Apoptotic Cells/ Unstimulated
SSC
FSC
Lymphocytes
FSC-A
FSC-H
Single Cells
Live-Dead UV
CD45-PepCpCy5.5
CD45 live
C11c-PECy7
BDCA1-BV605
BDCA1 DCs
pHrodo +
pHrodo-Red
SCARF1-APC
SCARF1+
F.
Apoptotic Cells
SSC
FSC
Lymphocytes
FSC-A
FSC-H
Single Cells
Live-Dead UV
CD45-PepCpCy5.5
CD45 live
C11c-PECy7
BDCA1-BV605
BDCA1 DCs
pHrodo +
pHrodo-Red
SCARF1+
SCARF1-APC
G.
Healthy Idividuals
% of Max
pHrodo +
Phagocytosis: pHrodo-AC
Full Serum
Ig Depleted Serum
H.
SLE Patients
% of Max
pHrodo +
Phagocytosis: pHrodo-AC

**Supplemental Figure 3. IgG depletion restores efferocytosis.** Ablation of IgG increases apoptotic cell uptake in SLE patients. (A) Diagram of experimental design. (B-D) IgG and autoantibodies to SCARF1 depleted from serum. We depleted 20% serum in RPMI, healthy or SLE, using 100μL of Protein A/G agarose beads columns. IgG depletion was confirmed by Coomasie stain (B) and Western Blot (C). SCARF1 binding was analyzed by Dot Blot. Recombinant protein was transfer to nitrocellulose membrane, and full or depleted serum was used as a primary antibody. Human IgG was used as secondary antibody. Representative blot (n=13 SLE and n=8 Healthy). Anti-SCARF1 was use a control. (E-H) Ig-Depletion restores efferocytosis in SLE patient serum. PBMCs ($1x10^6$/mL) were incubated with full serum, or Ig-depleted serum for 20–24 hours. FBS was used as serum control. (E-F) Flow cytometry schematic. Cells were incubated for 3 hours with pHrodo red-labeled apoptotic cells ($2x10^6$/mL). Cells were stained and analyzed by flow cytometry. (G-H) Representative histograms of pHrodo expression to measure phagocytosis. Blue line, full serum; Red line, Ig-depleted serum. Total number of $CD11c^+BDCA1^+SCARF1^+$ measure by flow cytometry in the presence of full serum or Ig-depleted serum. Data represent the mean (±SEM) of 2 independent experiments n=13 SLE and n=8 Healthy, by Two-way ANOVA.
